# Identification and Characterization of Neoplastic Cells in Keratin-Positive Giant Cell-Rich Tumors

**DOI:** 10.64898/2026.04.03.716202

**Authors:** M van der Linde, JSA Chrisinger, EG Demicco, CA Dehner, GW Charville, IH Briaire-de Bruijn, S Varma, C Zhu, M Matusiak, JVMG Bovée, M van de Rijn, DGP van IJzendoorn

## Abstract

Keratin-positive giant cell-rich tumor (KPGCT) is a newly described bone and soft tissue tumor. The tumor is characterized by scattered keratin-positive cells and the presence of *HMGA2::NCOR2* fusions. It is not known if the *HMGA2::NCOR2* fusion is located in the keratin-positive cells, and little is known about how KPGCT develops. KPGCT shares some histologic features with tenosynovial giant cell tumor (TGCT), a soft tissue tumor with *CSF1* rearrangements. Single-nuclei RNA sequencing (snRNA-seq) and Xenium spatial transcriptomics were used to elucidate the mechanisms driving KPGCT and compare KPGCT to TGCT. We show that the neoplastic cells in KPGCT constitute only a minority of cells in the tumor, and that they co-express keratin, *HMGA2* and *CSF1*. The neoplastic cells in KPGCT express no synovial markers, confirming KPGCT as a distinct entity, separate from TGCT. The bulk of the tumor consists of *CSF1R*-expressing macrophages and osteoclast-like giant cells, suggesting an important role for CSF1-CSF1R signaling. In addition, we find that the cells with the *HMGA2* translocation show activation of the hippo signaling pathway, which is known to regulate *CSF1* expression. We show that the CSF1-CSF1R axis, possibly regulated through the hippo signaling pathway, plays an important role in KPGCT. This axis likely stimulates the migration and proliferation of macrophages, which form the majority of cells in the tumor, as well as their differentiation into osteoclasts-like giant cells. These results provide a rationale for the use of CSF1R inhibitors, which have already shown efficacy in TGCT, as a therapy for KPGCT.

**Significance:** Keratin-positive giant cell-rich tumor (KPGCT) is a rare, newly described soft tissue and bone tumor. By examining this tumour on a single-cell level, we confirm the identity of the neoplastic cells on a molecular level, showing these form a minority of cells in the tumor. We show that activation of the hippo pathway in the neoplastic cells is a likely driver of tumorigenesis. Additionally, we show the neoplastic cells produce large amounts of CSF1, attracting the macrophages that form the majority of cells in the tumor. This finding gives supporting evidence for anecdotal reports of response to CSF1 inhibitor therapy. Finally, we identify key differences between KPGCT and tenosynovial giant cell tumor, a tumor that shares histological features with KPGCT.

## Introduction

Keratin-positive giant cell-rich tumor (KPGCT) is a recently described, rare soft tissue tumor. Clinically, the tumor presents as a swelling in either soft tissue or bone. Histologically, KPGCT displays nodules and lobules, surrounded by lymphoid cells. Multiple cell populations are often found in the lesions. Osteoclast-like and Touton-type giant cells are present, as well as large amounts of macrophages, which sometimes present as large fields of foamy histiocytes. The lesions occur most often in female patients between the ages of 10 and 40, but can be found at all ages. The tumors are often treatable with surgery and prognosis is generally good. However, individual cases with a more aggressive course, including metastasis, have been documented and long-term follow-up data is lacking (1–3).

A small subset of cells in the tumor express keratins as determined by pan-keratin immunohistochemistry. It was proposed that these cells represent the neoplastic component, though this has not yet been proven (1, 2, 4, 5). Keratin positivity is uncommon in soft tissue tumors, and understanding how keratin is expressed may provide insight into the pathogenesis of KPGCT (6–9).

KPGCT carries a recurrent *HMGA2* fusion, in most cases partnering with *NCOR2*. Additionally, a single case with a fusion of *HMGA2* with *COL14A1* was recently described (2–5, 10). It has also been shown at the RNA level that *CSF1* is expressed by a small subset of cells in the tumor, though it is not clear if these cells are the same cells as the keratin-positive cells (11, 12). Additionally, treatment with the CSF1R-inhibiting tyrosine kinase inhibitors imatinib and pexidartinib are effective in case reports describing unresectable KPGCT, indicating that CSF1 plays a role in KPGCT pathogenesis (3, 13).

In this study, we focused on uncovering the pathogenesis of KPGCT, as well as identifying possible targets for therapy. Using snRNA-seq, the neoplastic population of KPGCT was identified and key pathways activated within the neoplastic cells were characterized. In addition, we analyzed the tumor microenvironment, and identified neoplastic cell driven CSF1 signaling, providing a rationale for CSF1 inhibition as a targeted therapy.

## Methods

### snRNA-seq data generation

Formalin-fixed paraffin-embedded tissue from five KPGCT cases (Supplemental Table 1) were used for snRNA-seq, with five TGCT cases as comparison. Nuclei were isolated per the 10x Genomics protocol for FFPE tissue sections (GC000632) and subsequently prepared for sequencing with the 10x Genomics Chromium Fixed RNA Profiling Reagent Kit for Multiplex Samples (GC000527). Raw snRNA-seq data was subsequently pre-processed using the Cell Ranger workflow (v8.0.1).

### snRNA-seq quality control and processing

Unfiltered matrices generated with the Cell Ranger workflow were analyzed using the Seurat package (v5.3.0) in R (v4.4.2) (14). Low quality nuclei were removed based on the amount of features and mitochondrial RNA. Droplets with fewer than 400 features were excluded, as they were considered empty droplets, while nuclei with more than 2,500 features were excluded, as they likely represented doublets. Nuclei with a high percentage of mitochondrial RNA were also excluded. Cut-off points for mitochondrial RNA were determined on a case-by-case basis. Data from the different tumors was merged in a single Seurat object. Expression data was normalized and scaled, followed by a principle component analysis (PCA). Subsequently a nearest neighbor graph was constructed, and nuclei were clustered. Clustering was visualized through UMAP dimensional reduction.

### Integration and clustering

Data from multiple tumors was integrated with anchor-based CCA integration, using the PCA as an original reduction. This integrated data was then used to re-run the nearest neighbor and clustering calculations, followed by UMAP reduction. Pre- and post-integration UMAP visualization was compared to assess integration. Clusters were identified based on differentially expressed genes. Gene expression was calculated using the FindAllMarkers function in Seurat. Differentially expressed genes with an adjusted p-value < 0.05 were used to manually annotate clusters based on literature and the panglaoDB single cell database (15). DESeq2 was used to compare pseudobulked RNA from neoplastic cells in different tumors (16).

### Spatial Transcriptomics

A tissue micro array (TMA) was constructed, including a single 2mm core of KPGCT. The TMA was sectioned and processed at the Stanford Genomics Core Facility. Cells with 25 reads or less were removed during quality control. Data was normalized using SCTransform, after which principal component and neighborhood analysis, clustering and dimensional reduction were performed as described above. Spatial niches were identified using BANKSY, with lambda = 0.8 and k_geom = 50 (17).

### Identification of cell-cell interactions

Cell-cell interactions were identified using the CellChat (v2.1) R package (18). Analysis was limited to Secreted signaling proteins (e.g. CSF1, IL6, VEGF) and Cell-cell contact proteins (e.g. Notch), excluding matrix signaling such as collagen-integrin interactions. Cell signaling patterns were identified at k=6.

### Gene set enrichment analysis

Gene set enrichment analysis (GSEA) was performed with multiple R packages. Enrichment of gene ontology biological processes (GOBP) and Kyoto Encyclopedia of Genes and Genomes (KEGG) gene sets were performed with the FGSEA package (1.19.2), using the built-in gene sets (19). GSEA of transcription factor gene sets was performed using the clusterProfiler package (v4.14.6), using the MSigDB transcription factor targets genesets. Upregulated pathways were visualized using the enrichplot (v1.26.6) and pathview (v1.46.0) packages (20, 21). Only pathways with an adjusted p-value < 0.05 were used for further analysis.

### Immunohistochemistry

Formalin-fixed paraffin-embedded (FFPE) tissue was sectioned at a thickness of 3μm, deparaffinated and rehydrated, after which heat induced epitome retrieval at a pH of 10 was performed. Sections were incubated overnight with primary antibodies at 4°C. Primary antibodies used were wide spectrum screening cytokeratin (1:8,000, Dako, #Z0622) and CSF1 (1:6,400, Merck Life Science N.V., Clone 2D10, #MABF191). Sections were then incubated with secondary antibodies and developed with a peroxidase/alkaline phosphatase staining kit (ImmPRESS Duet, #MP-7714) according to the manufacturer’s instructions. Slides were scanned using the Vectra 3 Scanner (Akoya Bioscences).

### Immunofluorescence

Formalin-fixed paraffin-embedded (FFPE) tissue was sectioned at a thickness of 3μm, deparaffinated and rehydrated, after which heat induced epitope retrieval at a pH of 6 (citrate) or 9 (EDTA) was performed. Sections were incubated overnight with primary antibodies at 4°C. Primary antibodies used were pancytokeratin (1:50, pH6, Cell Signaling Technology, clone KeratinAE1/AE3, #67306), ser73 phosphorylated C-Jun (1:200, pH 6 Cell Signaling Technology, clone D47G9, #3270), GFPT2 (1:500, pH9, Abcam, clone EPR19095, #ab190966), and YAP1 (1:1200, pH6 clone 63.7, Santa Cruz, #sc-101199). Sections were then incubated with Alexa488, 555, 647 or 657 linked anti-mouse or -rabbit antibodies, followed by DAPI. Slides were scanned using the Pannoramic Midi II Scanner (3DHistech) or Keyence Fluorescence Microscope (BZ-X; Keyence).

### Immunofluorescence Analysis

Scanned images were analyzed using ImageJ (v1.54p). Nuclei were identified with localized Otsu thresholding on the DAPI stain, followed by watershedding to separate closely packed nuclei. Cells were identified as positive or negative for cytoplasmic stains based on individually identified thresholds for each slide. The average stain intensity of each nucleus was identified by serial application of cytoplasmic and nuclear masks identified above.

## Results

### Keratin-positive neoplastic cells express HMGA2

A total of 27,675 nuclei from five cases were sequenced, with repeat analysis for two cases, in seven separate barcoded libraries (Supplemental Table 1). Clustering by UMAP revealed thirteen clusters (Fig 1A). Only a small cluster of neoplastic cells was identified (Fig 1B). These neoplastic cells were identified with multiple methods. First, a measure of total combined keratin expression was calculated by adding the combined counts of the genes encoding for cytokeratins 1, 2, 3, 4, 5, 6, 7, 8, 10, 14, 15, 16, 17, 18 and 19, all of which are stained for by the cytokeratin AE1/AE3 antibody, which is used as a diagnostic immunohistochemical marker for KPGCT. This ‘keratin score’ was elevated in a cluster of cells with expression patterns highly similar to that of fibroblasts (Fig 1C, D). Further examination of keratins expressed by these cells showed mRNA expression of a total of 18 keratins, including high expression of *KRT16*, -*14*, -*8*, -*10* and -*7* (Supplement Fig 1). Second, expression of *HMGA2* was examined, and found to be limited to the cluster of keratin-expressing fibroblast-like cells (Fig 1C). Finally, expression of *GFPT2*, a marker for neoplastic cells in the histologically similar TGCT, was found solely in this cluster (Supplement Fig 2). As such, this cluster was identified as being the neoplastic component of KPGCT.

**Figure 1:**
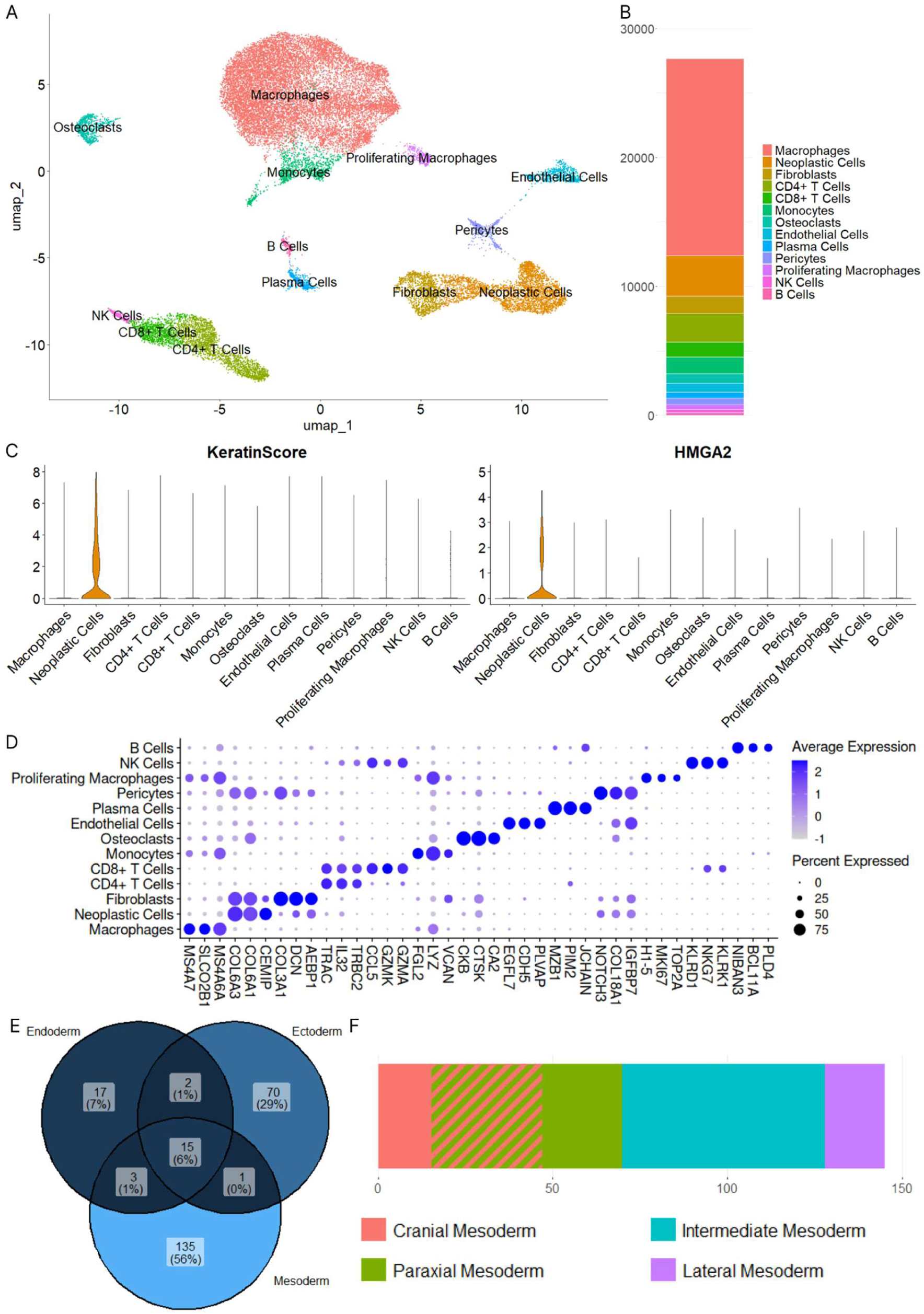
Overview of snRNA-seq data. **A)** UMAP plot of cells sequenced from 5 KPGCT cases. Large clusters of macrophages and lymphocytes were seen. A group of fibroblast-like *HMGA2* expressing cells was identified as the neoplastic population. **B)** Bar graph showing the amount of cells within each cluster in KPGCT. More than half of all cells are (proliferating) macrophages. Neoplastic cells form a minority of the cells in KPGCT. **C)** snRNA-seq expression of multiple keratins (left) and *HMGA2* (right) in KPGCT. Keratin and *HMGA2* expression is limited to neoplastic cells. **D)** Expression of the three most differentially expressed genes within each cluster. The neoplastic cells have an expression pattern similar to that of fibroblasts. Expression of proliferation markers is found within a subset of macrophages. **E)** Enrichment of developmental processes is concentrated on mesodermal processes. **F)** Enrichment of developmental processes within the mesoderm is concentrated on cranial/paraxial and intermediate mesodermal processes.

We further studied the phenotype of the neoplastic cells using a gene set enrichment analysis for terms in the GOBP database. We found the neoplastic cells were highly enriched for genes related to embryological development (262/440 gene sets). More detailed investigation showed enrichment for mesodermal processes, in particular of cranial, paraxial mesodermal or intermediate mesodermal development (Fig 1E, F). Within the intermediate mesodermal layer, processes related to kidney development were particularly common. We further studied the stages of embryonic development for expression of *HMGA2*, using the Descartes atlas (22). *HMGA2* was solely found within the kidney, specifically in the mesangial, metanephric and ureteric bud cells, which derive from the intermediate mesodermal layer (Supplemental Fig 3A). Additionally, expression was limited to early development (Supplemental Fig 3B), fitting the likely primitive nature of KPGCT.

### Macrophages constitute the majority of cells in the tumor

The identities of the remaining clusters were determined based on the expression of canonical marker genes. This resulted in the identification of macrophages, CD8+ and CD4+ T cells, NK cells, monocytes, B cells, plasma cells, pericytes, endothelial cells, osteoclast-like giant cells and a cluster of proliferating cells (Fig 1A, B, D, Supplement Fig 4, 5). Further analysis of the population of proliferating cells shows that these are a subset of macrophages, and not part of the keratin-positive neoplastic cells, similar to what has been reported in TGCT (23). Cell-cycle scoring by Seurat confirmed these cells were in the S-, G2- or M phase (Supplement Fig 6). While the neoplastic population represents only 12% of the cells, 55% of all cells are macrophages (Fig 1B), a subset of which expresses markers associated with proliferation. This observation is in line with histological and clinical descriptions of KPGCT as a slowly growing tumor, in which only a small proportion of cells are keratin-positive while macrophages form the bulk of tumor volume.

### KPGCT neoplastic cells express *CSF1*

CellChat was used to analyze the cell-cell communication network between the neoplastic cells and the tumor microenvironment, identifying ligand-receptor interactions. CellChat identified six distinct signaling patterns based on the involved ligand-receptor interactions, identifying similar patterns in both macrophages and proliferating macrophages, as well as different T cell subsets and NK cells (Supplement Fig 7). CellChat identified activation of the CSF1-CSF1R signaling pathway, in which the neoplastic cells interacted with the (proliferating) macrophages and osteoclast-like giant cells (Fig 2A, Supplement Fig 8). High expression of *CSF1* was found in the neoplastic population (Fig 2C). Additionally, there is expression of *CSF1R* in the proliferating macrophages, macrophages and osteoclast-like giant cells (Fig 2C). The expression of *CSF1* in the neoplastic cells was further confirmed with immunohistochemical co-staining for keratin and CSF1 (Fig 2B). Aside from *CSF1*, there was also a minor amount of *IL34,* a known ligand of CSF1R, and *IL6* expression in the neoplastic cells (Supplement Fig 9). Expression of *TNFRSF11A* (RANK) was found in the osteoclast-like giant cell population. However, no measurable expression of *TNFSF11* (the gene encoding for RANKL, which is a key signaling protein in osteoclast formation) was seen in the neoplastic population (Supplement Fig 9) (24).

**Figure 2:**
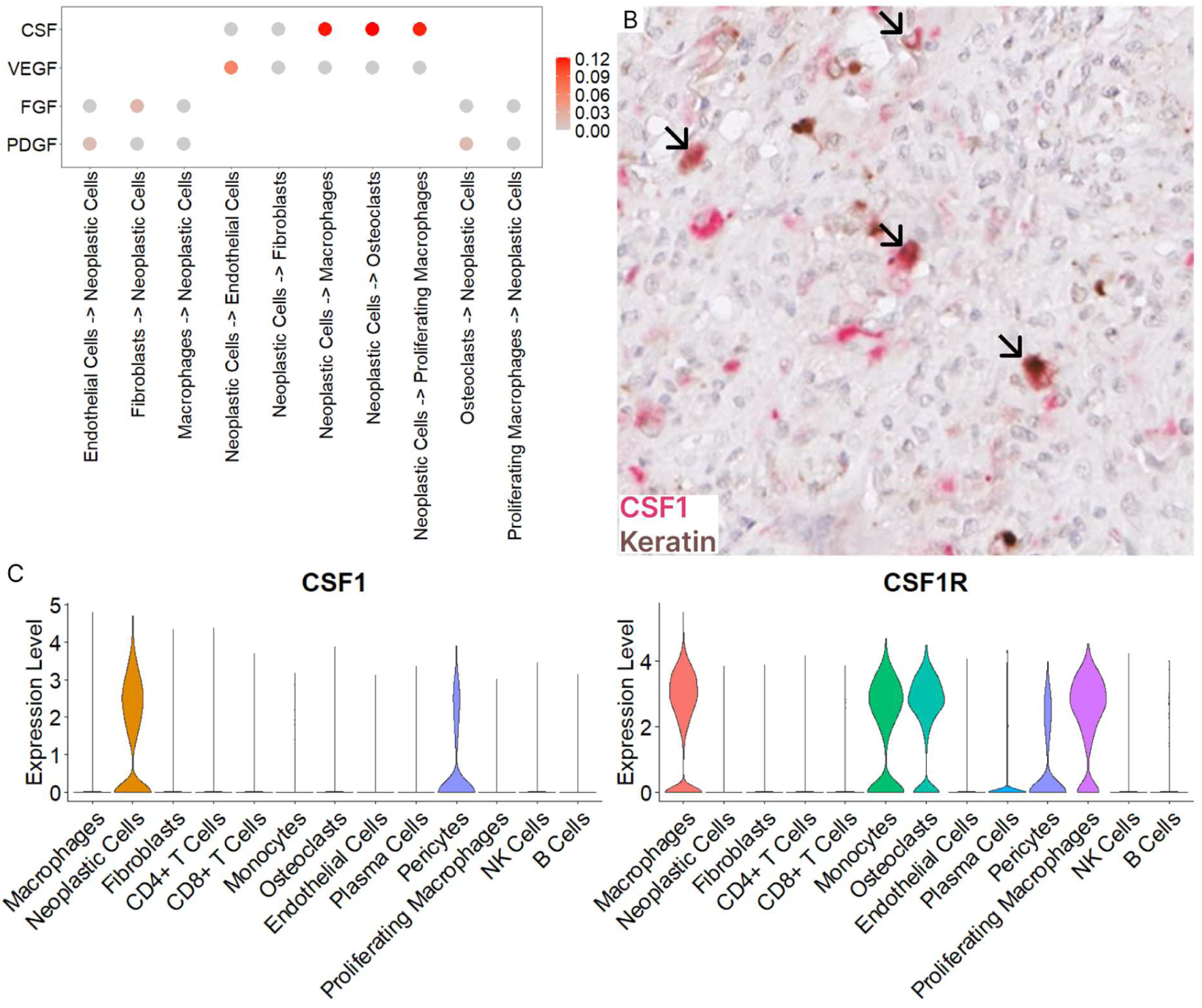
CSF1 signaling is a key pathway activated in KPGCT. **A)** Multiple different intercellular signaling pathways are active in KPGCT. A high level of CSF1 signaling is found between neoplastic cells and (proliferating) macrophages and osteoclast-like giant cells. Lower, but significant levels of VEGF, FGF and PDGF signaling are also present. **B)** IHC double stain in which CSF1 (pink) and keratin (brown) co-localize in neoplastic cells (Arrows), showing that the neoplastic cells of KPGCT produce CSF1 **C)** *CSF1* is highly expressed by neoplastic cells, while *CSF1R* is highly expressed by (proliferating) macrophages, monocytes and osteoclast-like giant cells, allowing for functional CSF1 signaling.

Three additional intercellular signaling pathways of note were identified by CellChat; the VEGF, FGF and PDGF signaling pathways (Fig 2A). First, VEGF signaling was identified between neoplastic cells and endothelial cells. Second, FGF signaling was identified between fibroblasts and neoplastic cells. Third, PDGF signaling was found from osteoclast-like giant cells and endothelial cells to neoplastic cells. Activity of these signaling pathways was confirmed by expression of the individual genes responsible for this signaling in the respective clusters (Supplement Fig 10).

### Hippo signaling pathway activation is found in KPGCT neoplastic cells

To examine which intracellular pathways are activated in the neoplastic cell population, driving the pathogenesis of KPGCT, a gene set enrichment analysis was performed. First, an enrichment analysis was performed using the KEGG database. This analysis yielded enrichments of multiple signaling-pathway related gene sets. Notably, there was a significant upregulation of the hippo pathway (Fig 3A). Further analysis of the genes within this pathway shows that there was increased expression of *CCN4*, a gene known to increase the activation of the hippo pathway (Supplement Fig 11). To assess to what extent the upregulation of genes within different intracellular signaling pathways resulted in changes in gene transcription, we performed gene set enrichment analysis for targets of transcription factors (i.e. genes transcribed by either RUNX2, AP1, TEAD1 or other transcription factors). This analysis yielded significant enrichment for multiple gene sets, most notably for two different gene sets which are commonly upregulated by the transcription factor TEAD1, a key transcription factor in the hippo pathway (WGGAATGY TEF1 Q6 and TEF1 Q6, Fig 3B).

**Figure 3:**
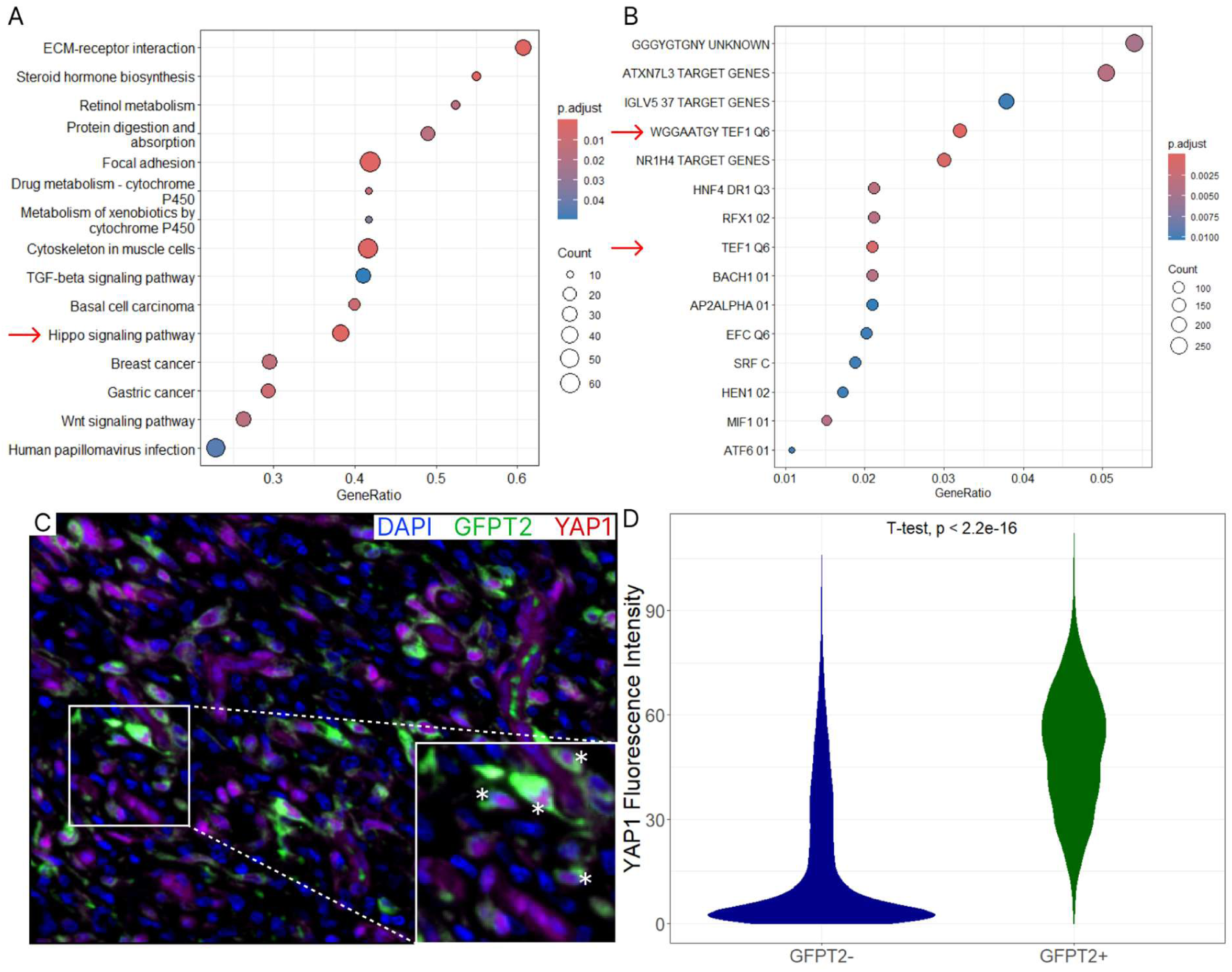
The hippo pathway is activated in KPGCT. **A)** Gene set enrichment analysis shows that genes in the hippo signaling pathway are highly expressed in KPGCT (‘Hippo signaling pathway’, red arrow). **B)** Gene set enrichment analysis shows that targets of the transcription factor TEAD1 are upregulated (‘WGGAATGY TEF1 Q6’ and ‘TEF1 Q6’, red arrows). **C)** Immunofluorescent staining for YAP1 and GFPT2 shows nuclear YAP1 in all GFPT2+ cells (asterisks), while GFPT2-cells have variable YAP1 staining. **D)** Quantification of nuclear YAP1 signal intensity shows a strong increase of nuclear YAP1 positivity in GFPT2+ cells (p < 0.05).

Activation of the hippo pathway was verified through YAP1 immunofluorescence. YAP1 immunofluorescence showed nuclear positivity in the neoplastic cells as identified by GFPT2 positivity (Fig 3C, white asterisk). In contrast, nuclear YAP1 positivity was only rarely seen in the non-neoplastic cells. YAP1 positivity in these cells was generally limited to the cytoplasm, as seen with inactive YAP1 (Fig 3C, white arrow). This result is confirmed by the quantification of the nuclear YAP1, in which we examined the signal intensity of YAP1 in GFPT2-positive and GFPT2-negative cells. A total of 7,920 GFPT2-positive and 38,830 GFPT2-negative cells were analyzed, showing a significant increase of nuclear YAP1 intensity in the GFPT2-positive neoplastic cells, as compared to the GFPT2-negative non-neoplastic cells (49.7 vs 16.1, p<0.05, see Fig 3D).

### JUN transcription factor is active in KPGCT neoplastic cells

To examine how expression of keratin is upregulated in KPGCT, we compared neoplastic KPGCT nuclei to nuclei from neoplastic cells of TGCT, a similar but keratin-negative tumor. A total of 84,953 TGCT nuclei were sequenced. Neoplastic cells were identified as fibroblast-like cells with elevated *CSF1* expression, and further verified with *GFPT2* and *CLU*, both known markers for neoplastic cells in TGCT (23). Remaining clusters were identified based on canonical marker genes. Integration of KPGCT and TGCT datasets shows a significant overlap between the non-neoplastic components of both tumors (Fig 4A). In contrast, the neoplastic cells of KPGCT and TGCT formed neighboring subclusters within the neoplastic cell cluster (Fig 4A).

**Figure 4:**
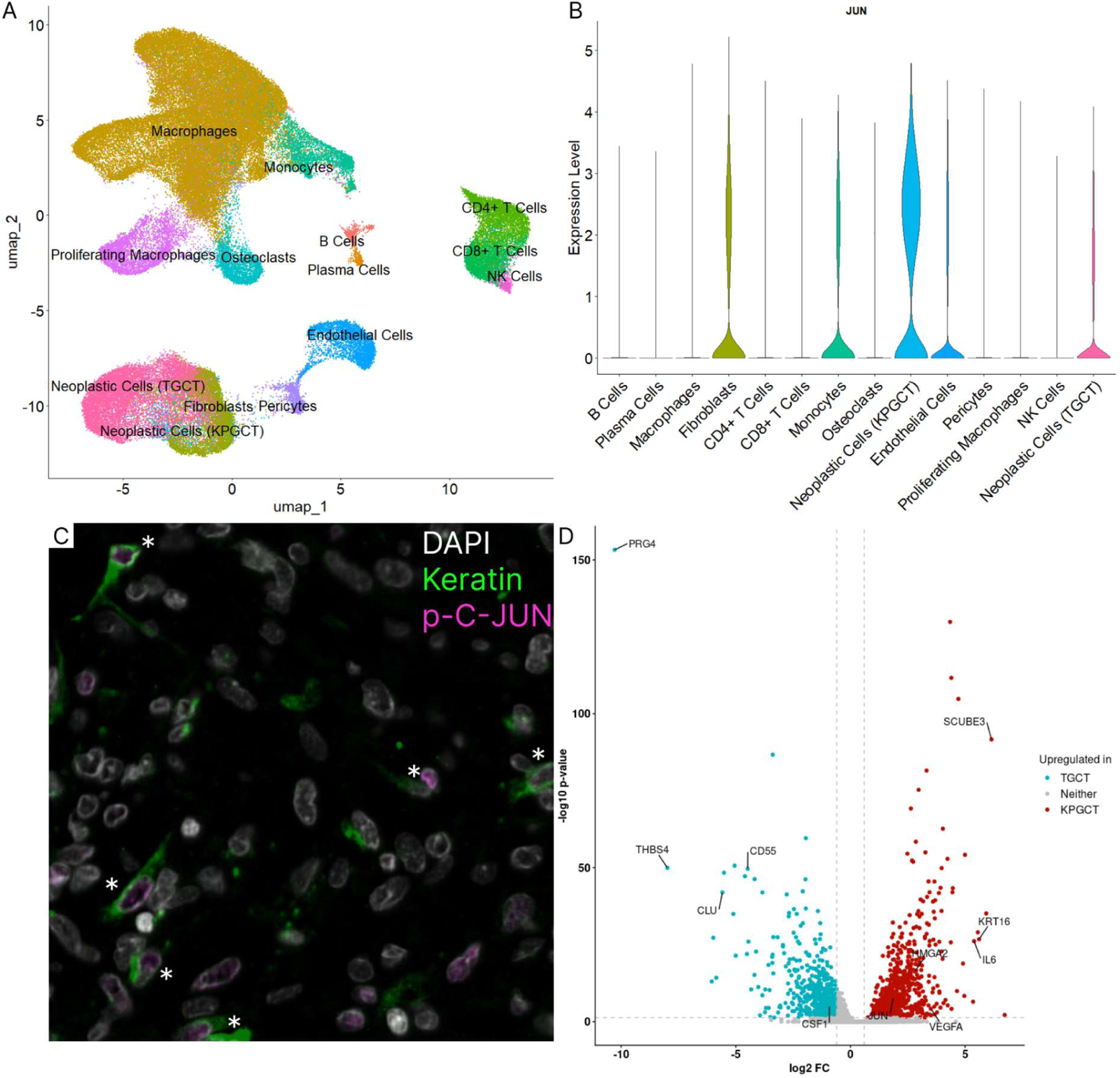
KPGCT is similar, though not identical, to TGCT. **A)** UMAP plot of cells from 5 KPGCT and 5 TGCT cases, forming distinct subclusters within the group of neoplastic cells. **B)** snRNA-seq expression of *JUN*, showing high expression in KPGCT. **C)** Immunofluorescence stain of KPGCT, stained for keratin and phosphorylated C-JUN. Phosphorylated C-JUN is found in keratin+ cells(*), while being nearly absent in other cells. **D)** Differential gene expression of TGCT versus KPGCT highlights increased expression of *HMGA2*, *JUN, KRT16* and multiple signaling molecules in KPGCT as compared to TGCT. Increased expression of *PRG*, *CLU, THBS4* and *CSF1* is found in TGCT.

Many of the highly expressed keratins in the neoplastic cells of KPGCT are known to be regulated by C-JUN (25, 26). Examination of *JUN* expression in KPGCT and TGCT shows that *JUN* is highly expressed in KPGCT neoplastic cells, while expression is significantly lower in TGCT neoplastic cells (Fig 4B). Additionally, gene set enrichment analysis of transcription factor targets showed increased activity of the AP-1 transcription factor complex, of which C-JUN is a key component. To verify activity of C-JUN, we performed immunofluorescence staining for S73 phosphorylated C-JUN. This showed nuclear positivity in keratin-positive cells, confirming activation of C-JUN (Fig 4C).

Integrated data from the neoplastic cells in the five cases of KPGCT and TGCT was used to identify similarities and differences between the neoplastic cell populations of both tumors. When examining the differentially expressed genes in TGCT and KPGCT neoplastic cells, about one third of genes were expressed in both tumors, while the remaining genes were only expressed in one of the two tumors (Supplemental Fig 12). Increased expression of *KRT16, HMGA2, JUN and SCUBE3* were found in KPGCT neoplastic cells, while *CLU, CD55, THBS4 and PRG4*, each proposed markers for synoviocytes or TGCT neoplastic cells (23, 27–29), were upregulated in TGCT neoplastic cells (Fig 4D, Supplemental Data 1). Additionally, the level of *CSF1* expression was found to be higher in TGCT neoplastic cells, while *VEGFA* and *IL6* were expressed higher in KPGCT neoplastic cells.

### Xenium spatial transcriptomics confirms activation of CSF1 and hippo signaling in KPGCT

To verify the signaling pathways identified with snRNA-seq, Xenium spatial transcriptomics was performed on KPGCT3, and cell clusters annotated by markers identified in snRNA-seq (Supplemental Fig 13A). The upregulation of *CCN4* in neoplastic cells was confirmed by Xenium spatial transcriptomics (Fig 5A). Expression of *CSF1* was found in neoplastic cells, and *CSF1R* expression was seen in macrophages, osteoclast-like giant cells, and other myeloid cells (Fig 5B). To confirm the chemoattractive effects of neoplastic cells on the macrophage population, the distance was calculated between the centroids of each macrophage and the closest neoplastic cell, as well as the distance to the closest non-macrophage, non-neoplastic cell. This distance calculation revealed that macrophages tend to locate near neoplastic cells (mean distance to nearest neoplastic cell = 11.72µm, mean distance to nearest other cell = 14.00µm, p < 0.05), providing additional evidence of chemoattractive factors being produced by neoplastic cells (Fig 5C).

**Figure 5:**
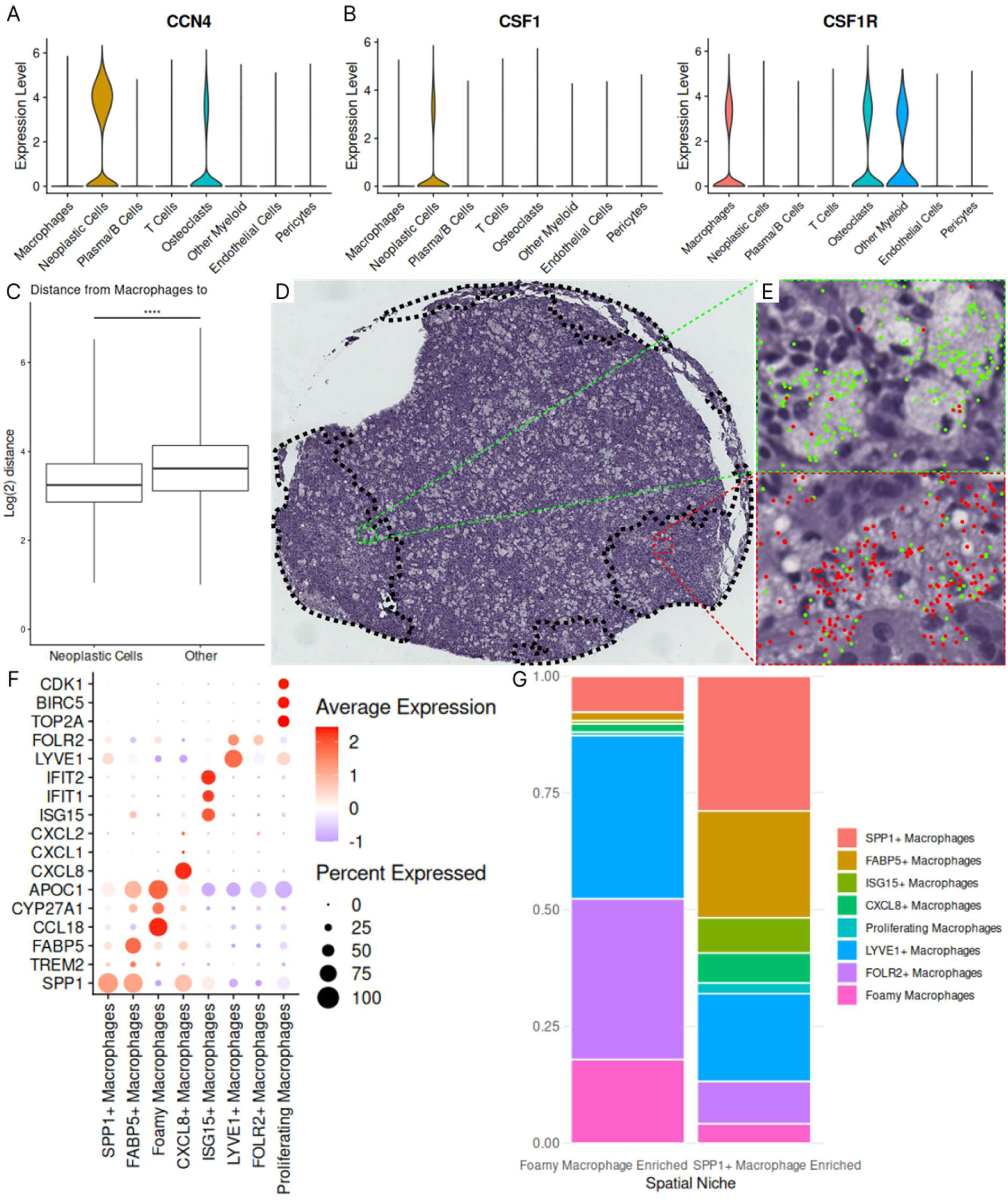
Xenium spatial transcriptomics shows spatial organization in KPGCT. **A)** *CCN4* expression is seen in neoplastic cells, confirming the *CCN4* driven activation of the hippo pathway in neoplastic cells. **B)** *CSF1* expression is found in neoplastic cells. *CSF1R* is found in (proliferating) macrophages and other myeloid cells, confirming the neoplastic cell driven proliferation of macrophages. **C)** The distance from macrophages to the nearest neoplastic cell and nearest other, non-neoplastic, cell. Macrophages localize closer to the neoplastic cells, in line with the chemoattractive effects of CSF1 signaling. Boxplot shows 50% and 90% intervals. ****P<0.00005 **D)** BANKSY identified niches overlaid on the tissue core used for Xenium. A central area enriched with foamy cells and peripheral areas poor in foamy cells (SPP1+ macrophage niche) are seen. **E)** Expression of *CYP27A1* and *CCL18* (green), is higher in large foamy macrophages (top), while expression of *TREM2* and *FABP5* (red) is higher in small foamy macrophages (bottom). **F)** Markers for macrophage subtypes. **G)** Distribution of macrophage subtypes in the foamy macrophage-enriched niche and the SPP1+ macrophage-enriched niche.

### The KPGCT microenvironment is divided into distinct niches

To analyze the KPGCT microenvironment, we applied the BANKSY algorithm to identify spatial niches. Two main niches were identified, however the distribution of general cell types was identical between the different spatial niches (Supplemental Fig 13B). Therefore, we examined macrophage subtypes as a possible factor influencing niche formation, using both the histology and Xenium morphology images (Fig 5D). These showed both foamy macrophages (Supplemental Fig 14, asterisks) and multinucleated giant cells (Supplemental Fig 14, arrows), with localized increases in foamy macrophages, leading us to suspect that macrophage subtypes characterize different spatial niches.

To better understand the macrophage subtypes characterizing different spatial niches, we sub-clustered of macrophages, and identified distinct subsets of macrophages, consisting of SPP1+, FABP5+, ISG15+, CXCL8+, LYVE1+, FOLR2+, proliferating and foamy macrophage subtypes (Fig 5E, F) (30). The presence of these macrophage subtypes was confirmed through analysis of macrophages in the snRNA-seq data generated earlier (Supplemental Fig 15). In particular, we compared cell morphology and gene expression, resulting in the identification of multiple markers for large foamy macrophages, including *CCL18* and *CYP27A1.* Expression of *TREM2* and *FABP5* was low in these cells, and was found primarily in a subset of SPP1+ macrophages. Mapping these cells back to the segmentation stains and H&E stain showed that *TREM2* and *FABP5* were expressed in small foamy macrophages, rather than large foamy macrophages (Fig 5E). Expression of *APOC1* was found in both macrophage subtypes (Fig 5F).

The identified macrophage subtypes were mapped back to the two identified niches, showing enrichment of SPP1+ macrophages in one niche, as well as increases in interferon stimulated and CXCL8+ macrophages. The other niche was characterized by an increase in foamy macrophages, as well as FOLR2+ macrophages. There was no significant difference in the distribution of the LYVE1+ or proliferating macrophages between the two niches (Fig 5G). Examination of the spatial distribution of these macrophage subtypes showed that those found in the SPP1+ macrophage niche tended to colocalize in smaller, distinct niches, while those found in the foamy macrophage niche were more intermingled with each other (Supplemental Fig 16).

## Discussion

In this study we aimed to identify and characterize the neoplastic and non-neoplastic cells in KPGCT using snRNA-seq and Xenium spatial transcriptomics. We identified the KPGCT neoplastic cells and show these cells are keratin-positive. The probe-based snRNA-seq technology required for FFPE material does not allow for the identification of DNA fusions, such as the characteristic *HMGA2::NCOR2* fusion found in KPGCT. However, *HMGA2* expression was limited to a well-defined cluster of keratin-positive cells. As *HMGA2* expression is both common in mesenchymal neoplasms and absent in most adult tissue (31, 32), this shows keratin-positive cells are the neoplastic component.

While KPGCT neoplastic cells express multiple keratins, gene set enrichment analysis confirms they have a mesenchymal origin. Expression of *JUN* is found in KPGCT, while being absent in TGCT. C-JUN is a transcription factor known to regulate the expression of multiple keratins (26), many of which are expressed by KPGCT neoplastic cells. The expression of *JUN* could explain why KPGCT is keratin-positive.

Notably, while both KPGCT and TGCT show similar histology and hippo pathway activation, expression of synovial markers such as *CLU* and *PRG4* (also known as lubricin, a key protein for lubrication of the synovial space) is absent in KPGCT (27, 29). This finding indicates that KPGCT does not arise from the synovium, and should be regarded as a distinct entity. Recently, keratin negative KPGCT have been reported, posing a challenge for pathologists in differentiating these from TGCT in the absence of molecular testing (33). Additionally, we have shown that KPGCT is positive for CSF1 immunohistochemistry, further compounding this problem, as CSF1 immunohistochemistry or chromogenic in situ hybridization can be used to identify TGCT when histology alone is insufficient (34, 35). In this context clusterin immunohistochemistry, which may be used to identify TGCT neoplastic cells (27), could be a useful diagnostic tool to distinguish TGCT from (keratin negative) KPGCT.

As expected, we found that macrophages and osteoclast-like giant cells form the majority of cells in the KPGCT tumor mass, confirming prior histological descriptions (1, 2, 4, 5). Single cell and spatial transcriptomics analyses in the current study provided further insight into the nature of these macrophages. Notably, the only proliferating cells identified were macrophages. A high level of *CSF1* expression is seen in KPGCT neoplastic cells, confirming earlier studies demonstrating *CSF1* expression in KPGCT (11, 12, 36). Additionally, a high level of *CSF1R* expression is present in both macrophages and osteoclast-like giant cells, but not in neoplastic cells, suggesting a predominately paracrine rather than autocrine role for CSF1 secretion in these tumors, similar to what is seen in TGCT (23). This hypothesis is supported by the spatial distribution of cells shown by Xenium spatial transcriptomics, wherein macrophages are located close to neoplastic cells. CSF1-CSF1R signaling attracts macrophages, and is essential for macrophage proliferation and their differentiation into osteoclast-like giant cells (37–40). Hence, it is likely that the activation of the CSF1-CSF1R signaling pathway explains both the macrophage and giant cell-rich histology of KPGCT. Therapeutic antibodies targeting CSF1 and CSF1R, as well as small molecule inhibitors of CSF1R have been developed, and are effective in TGCT, a similar CSF1-driven giant cell-rich tumor (41–45). Additionally, case reports describe a good response to CSF1 inhibition in irresectable KPGCT (3, 13). Our findings provide a rationale for the use of these treatments in KPGCT.

Conversely, there was no expression of *TNFSF11* (RANKL), an important regulator of osteoclast formation, which is thought to cause giant cell formation in giant cell tumor of bone (24, 46). This is consistent with case reports in which KPGCT was treated with the RANKL inhibitor denosumab, with limited clinical effect (2, 12).

We show an increased expression of *CCN4*, which is known to inhibit YAP1 degradation (47), as well as an increase in expression of genes transcribed by the TEAD1 transcription factor, consistent with activation of the hippo pathway (48). Finally, we confirm activation of the hippo pathway in KPGCT using YAP1 immunofluorescence. Notably, activation of the hippo pathway is also seen in TGCT (23). The activation of the hippo pathway may offer a novel therapeutic target in both lesions. While no TEAD inhibitors have been tested in phase 3 trials, a phase 1/2 trial using the YAP/TEAD inhibitor VT3989 in tumors with persistent TEAD activation (mesothelioma, epithelioid hemangioendothelioma and meningioma) has shown marked clinical benefit (49). While further research is necessary, this would allow for direct targeting of neoplastic cells in KPGCT, and may be more efficient than targeting the CSF1-CSF1R axis.

HMGA2 fusions may be found in multiple soft tissue tumors, as well as some epithelial lesions. The exact mechanism by which these fusions cause tumors to develop is unknown and different fusion partners are found in different lesions, each appearing to activate distinct mechanisms in different tumors (32, 50–55). While it seems likely that the *HMGA2* fusion causes activation of the hippo pathway, it is not clear if this is the result of *HMGA2* overexpression or a possible gain of additional functions due to the *HMGA2::NCOR2* fusion. Little is known about the functioning of HMGA2 in general, and more research is needed to elucidate how the *HMGA2::NCOR2* fusion causes KPGCT formation.

To summarize, we have used snRNA-seq to examine possible mechanisms of pathogenesis in KPGCT, and signaling networks between neoplastic cells and their microenvironment. We show that the neoplastic cells of KPGCT express *HMGA2* and multiple keratins. Additionally, we show that the neoplastic cells produce CSF1 and interact with macrophages, likely attracting them and inducing proliferation and differentiation into osteoclast-like giant cells. This signaling axis is an attractive therapeutic target, for which therapeutic antibodies and small molecule inhibitors exist. Our data suggest that the expression of *CSF1* in KPGCT may be driven by activation of the hippo pathway, while C-JUN activation may explain the expression of keratins. Finally, we show that KPGCT likely does not arise from synovium, and is a distinct entity, separate from TGCT. We also show similarities and differences between KPGCT and TGCT, and provide possible markers for distinguishing these tumors.

## Data, Materials, and Software Availability

Raw sequencing data has been uploaded the National Center for Biotechnology Information (NCBI) Gene Expression Omnibus (GSE326793)

## Author Contributions

**MvdL**: Investigation, Formal analysis, Experimental validation, Visualization, Writing, Editing. **JSAC, EGD, CAD, GWC, MM**: Resources, Editing. **IHBB**, **SV**: Experimental validation. **CZ**: Data generation. **JVMGB**: Resources, Editing, Supervision. **MvdR**: Resources, Editing, Conceptualization, Supervision. **DGPvIJ**: Review, Editing, Conceptualization, Supervision.

## Supplemental Tables

**Supplemental Table 1.** Clinico-pathological features of KPGCT cases used for snRNA-seq.

| Case | Age | Sex | Location | Size | Molecular Analysis | Keratin positive |
| --- | --- | --- | --- | --- | --- | --- |
| 1 | 36 | M | Ilium | 14 cm | *HMGA2* exon 3::<br>*NCOR2* exon 16 | Yes |
| 2 | 20 | F | Thigh | 4 cm | *HMGA2* exon 2::<br>*NCOR2* exon 16 | Yes |
| 3 | 15 | M | Anterior skull base and frontal lobe | 7.7 cm | *HMGA2* exon 3::<br>*NCOR2* exon 16 | NA |
| 4 | 87 | F | Tibia | 6.2 cm | *HMGA2* exon 3::<br>*NCOR2* exon 16 | NA |
| 5 | 20 | F | Subcutaneous soft tissue of back | 10 cm | *HMGA2* exon 3::<br>*NCOR2* exon 16 | Yes |

## Supplemental Figures

**Supplemental Figure 1.**
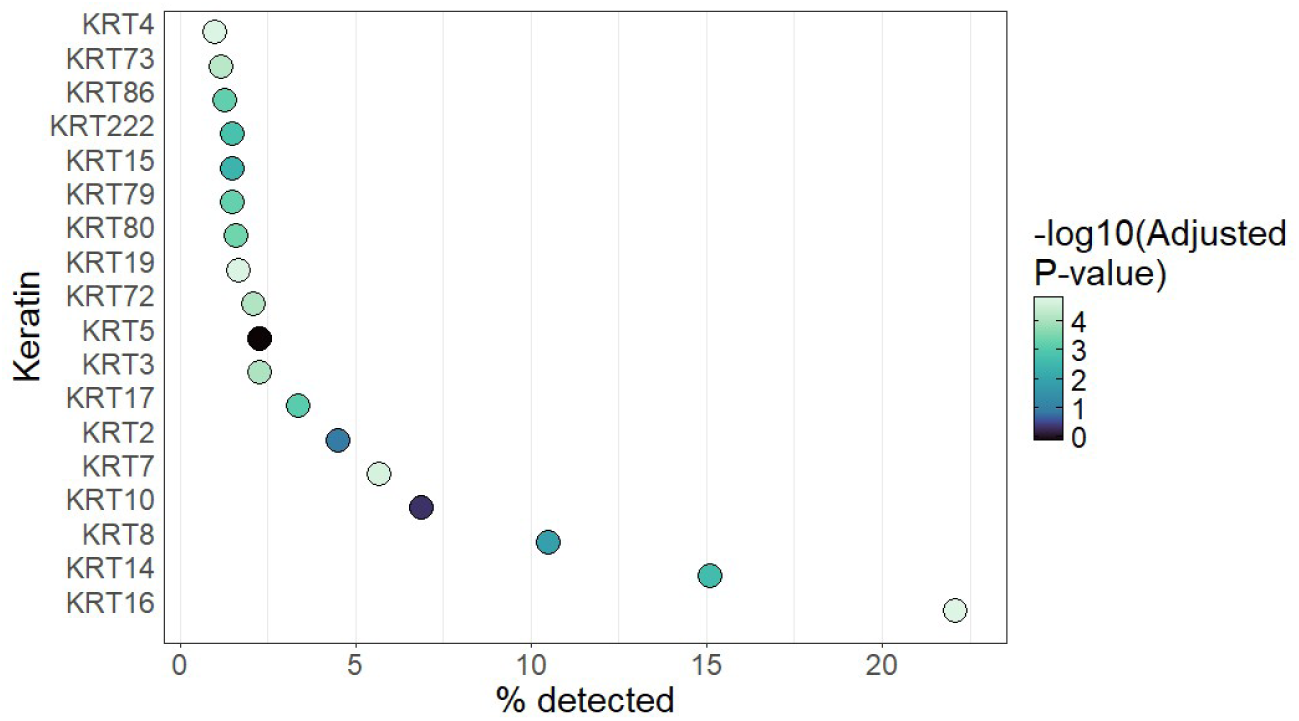
Expression of keratins within KPGCT neoplastic cells

**Supplemental Figure 2.**
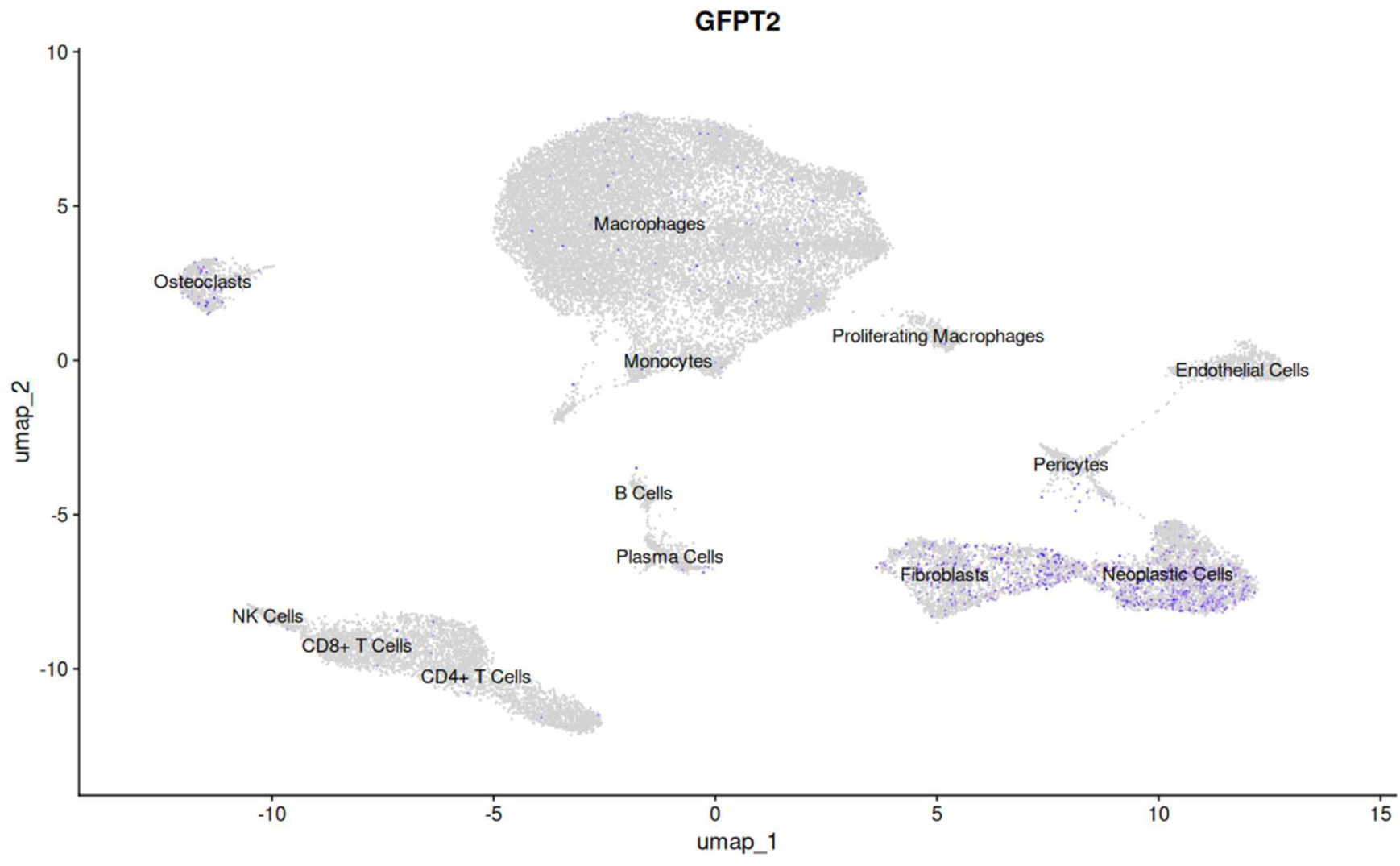
GFPT2 expression is seen in neoplastic cells

**Supplemental Figure 3.**
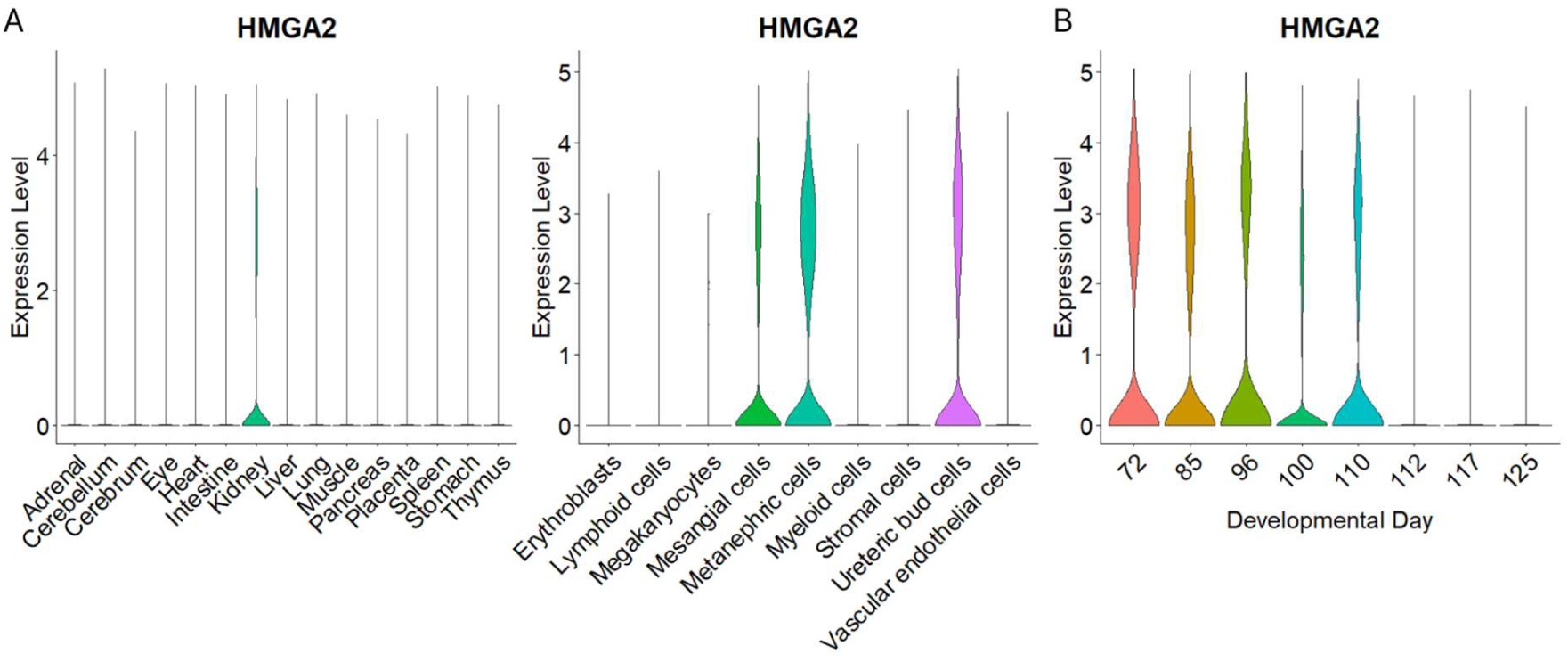
A) Expression of *HMGA2* in embryonic tissue is found solely within the kidney. B) Within the embryological kidney *HMGA2* expression is only found in early intermediate mesoderm-derived cells. Data from Cao et al.(**22**)

**Supplemental Figure 4.**
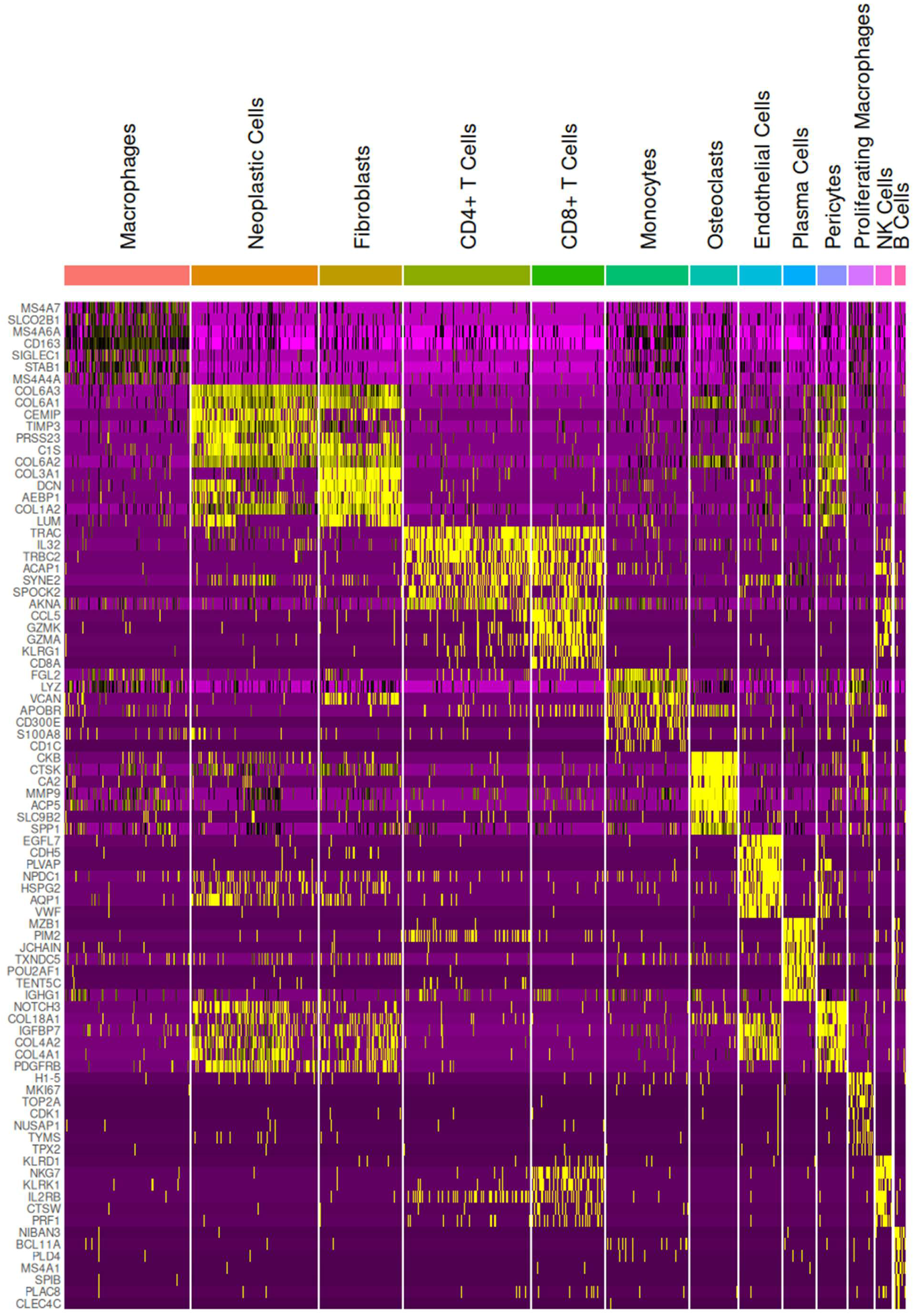
Expression of genes within the different cell types in KPGCT

**Supplemental Figure 5.**
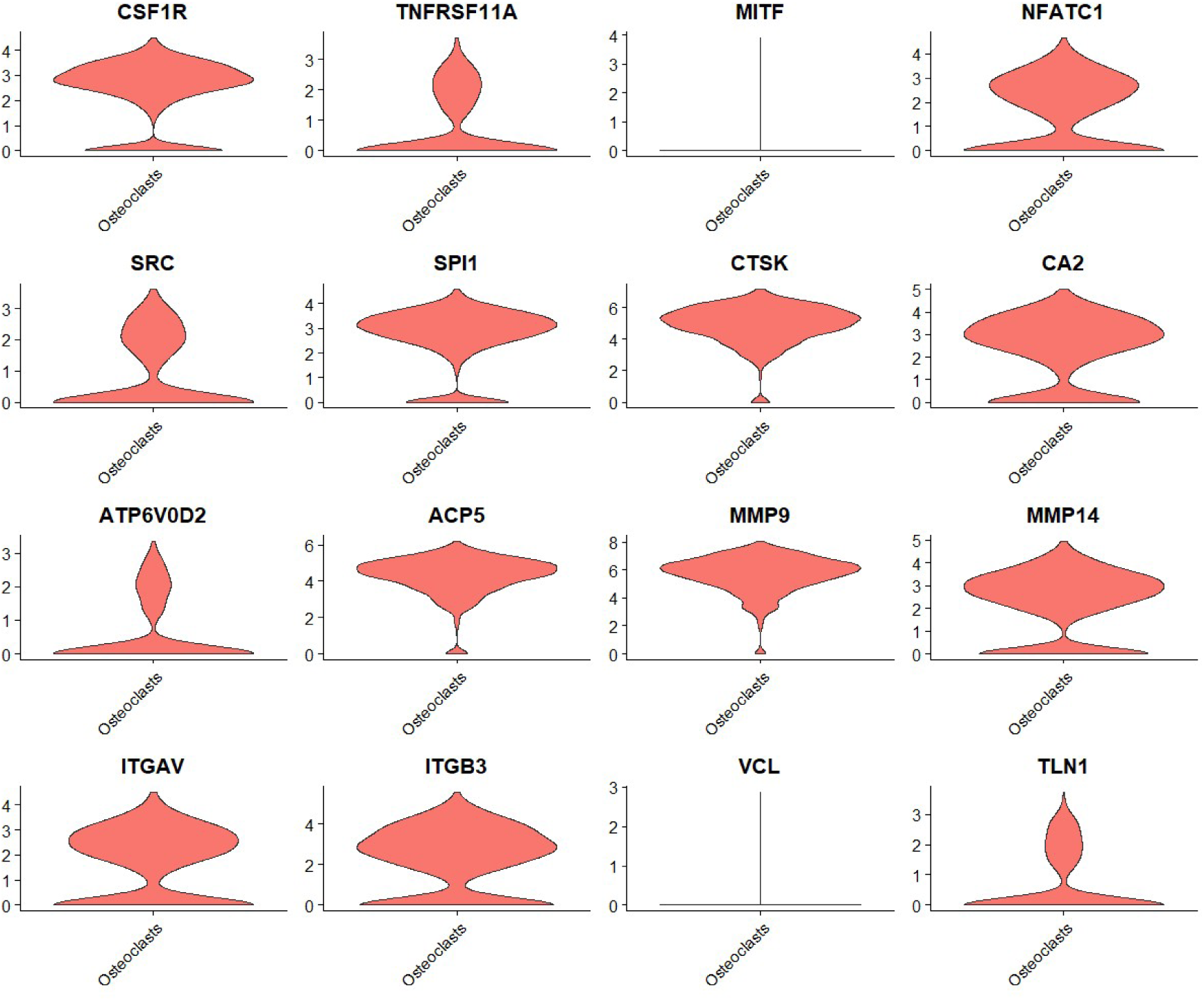
Expression of osteoclast genes within the ‘Osteoclast’ cell cluster

**Supplemental Figure 6.**
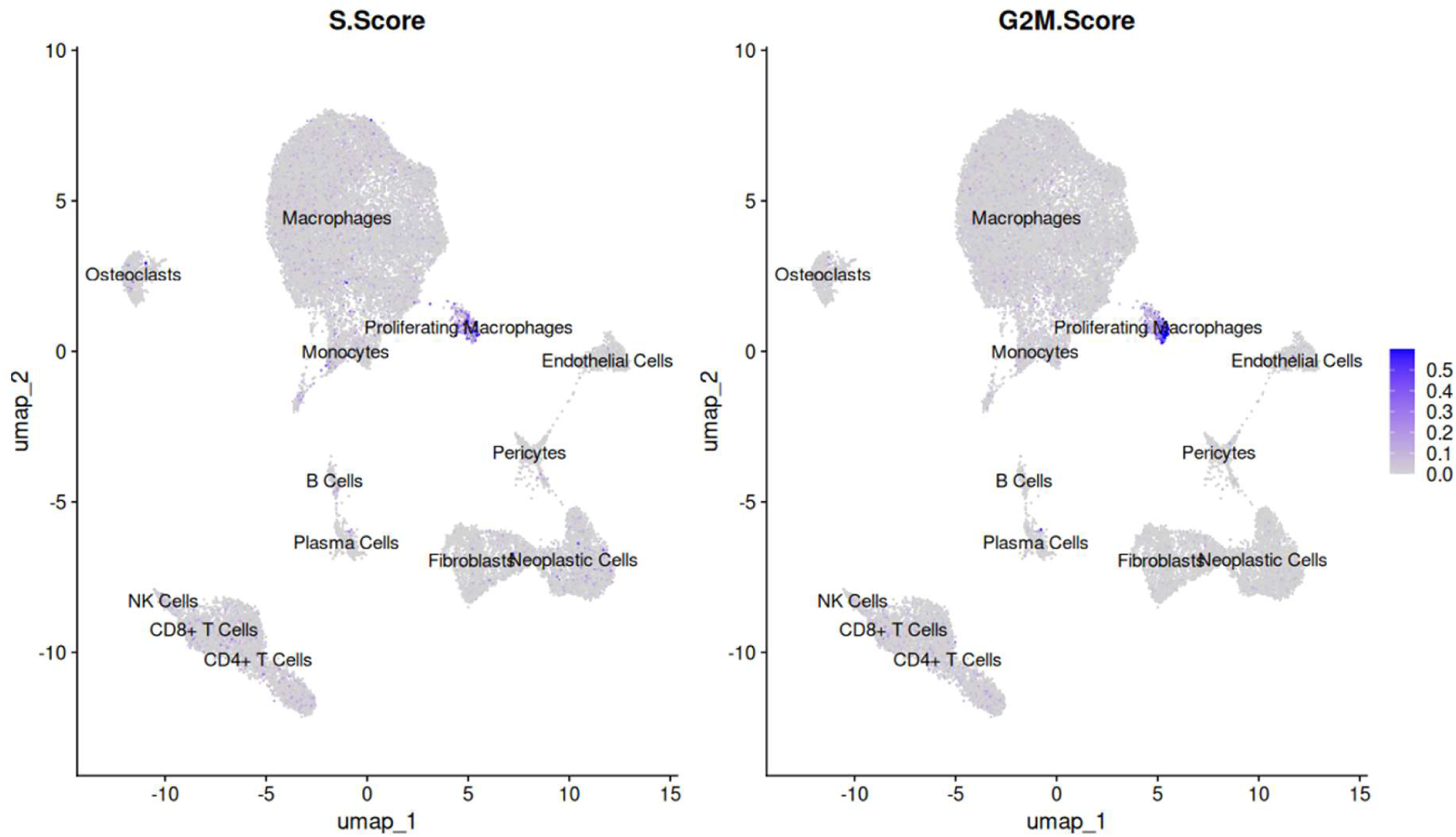
S- and G2M-phase related genes, expressed as S.Score and G2M.score is increased in a small cluster of macrophages

**Supplemental Figure 7.**
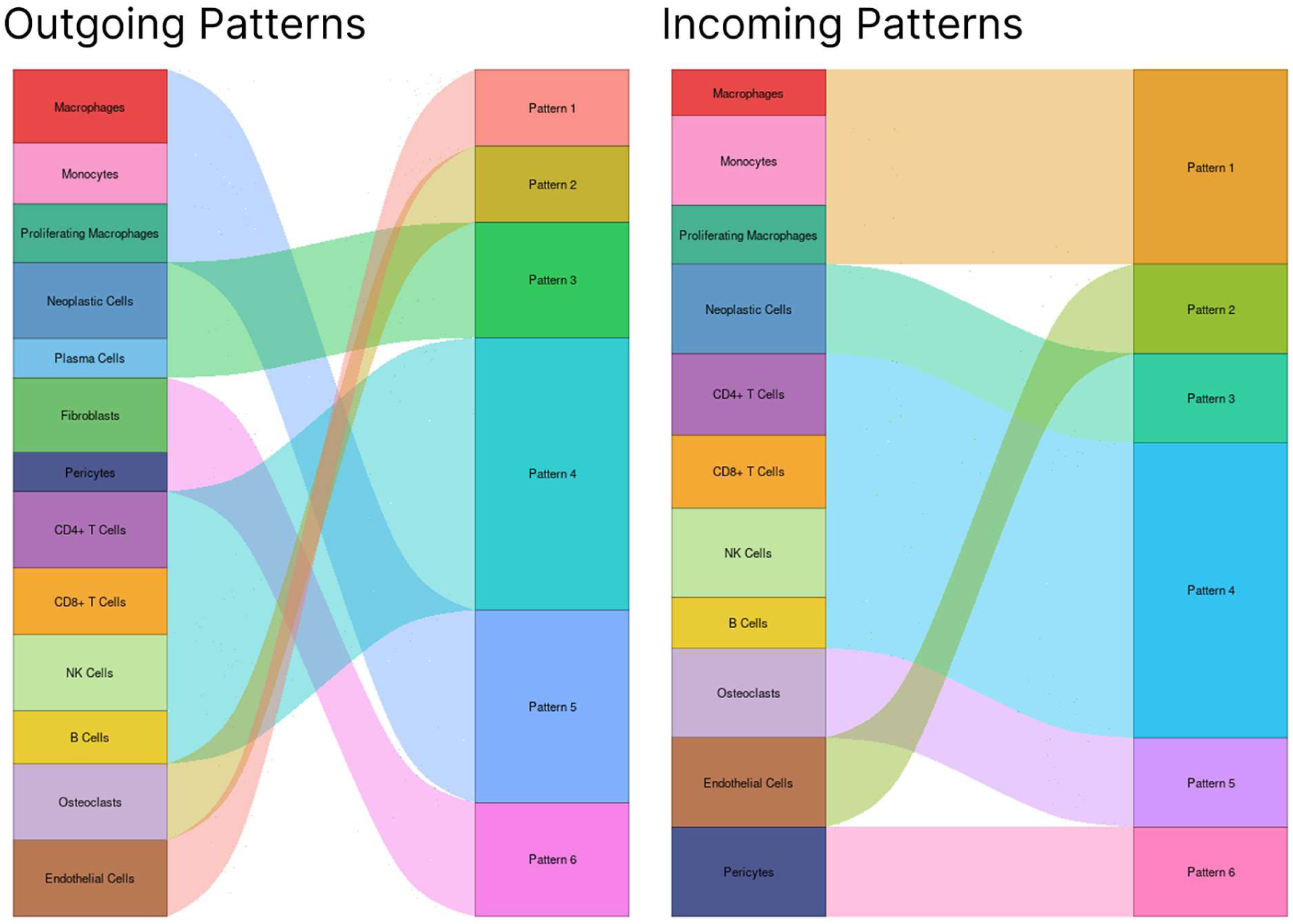
Patterns of cell signaling as identified by cellchat

**Supplemental Figure 8.**
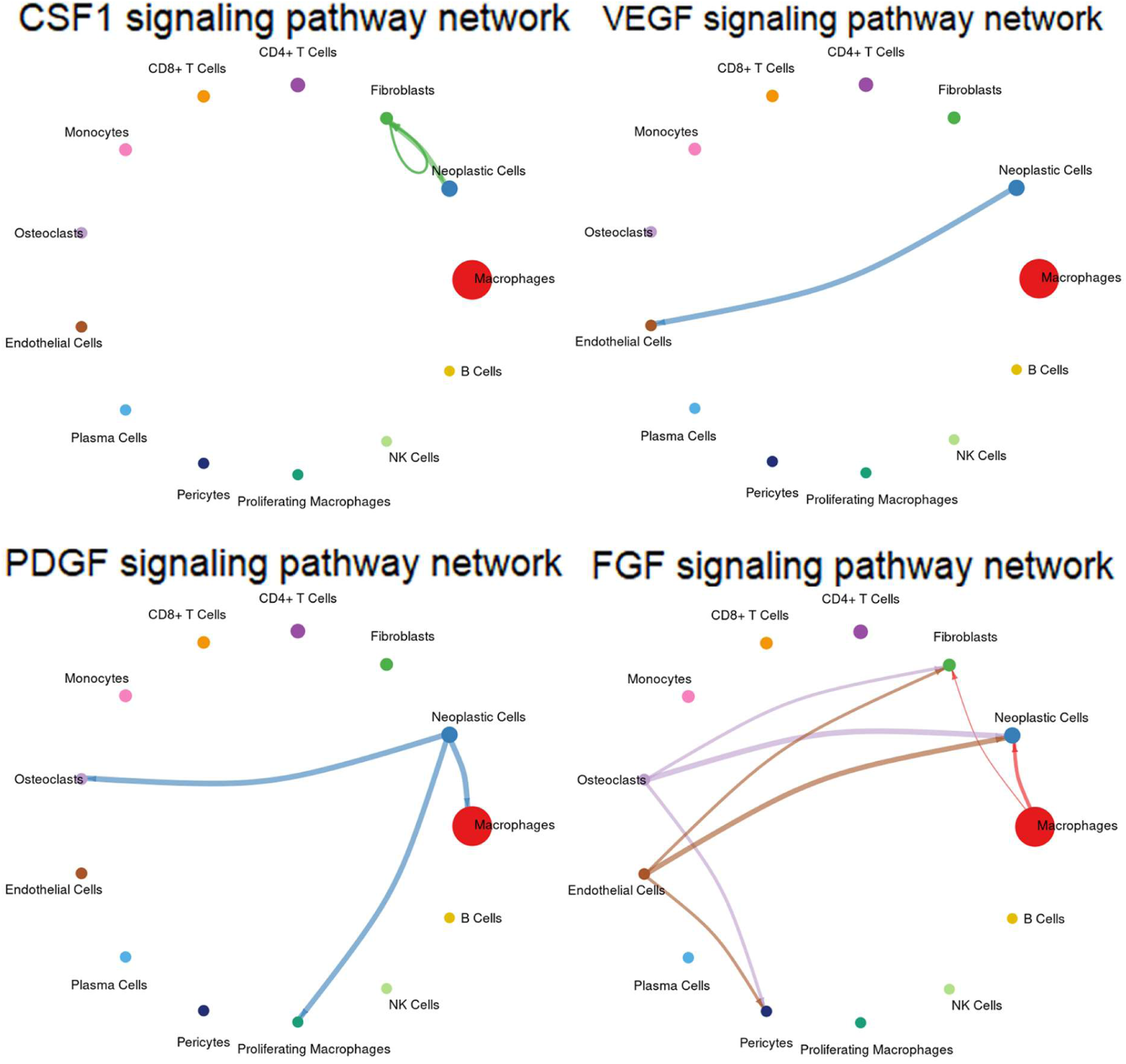
Signaling networks as identified by cellchat

**Supplemental Figure 9.**
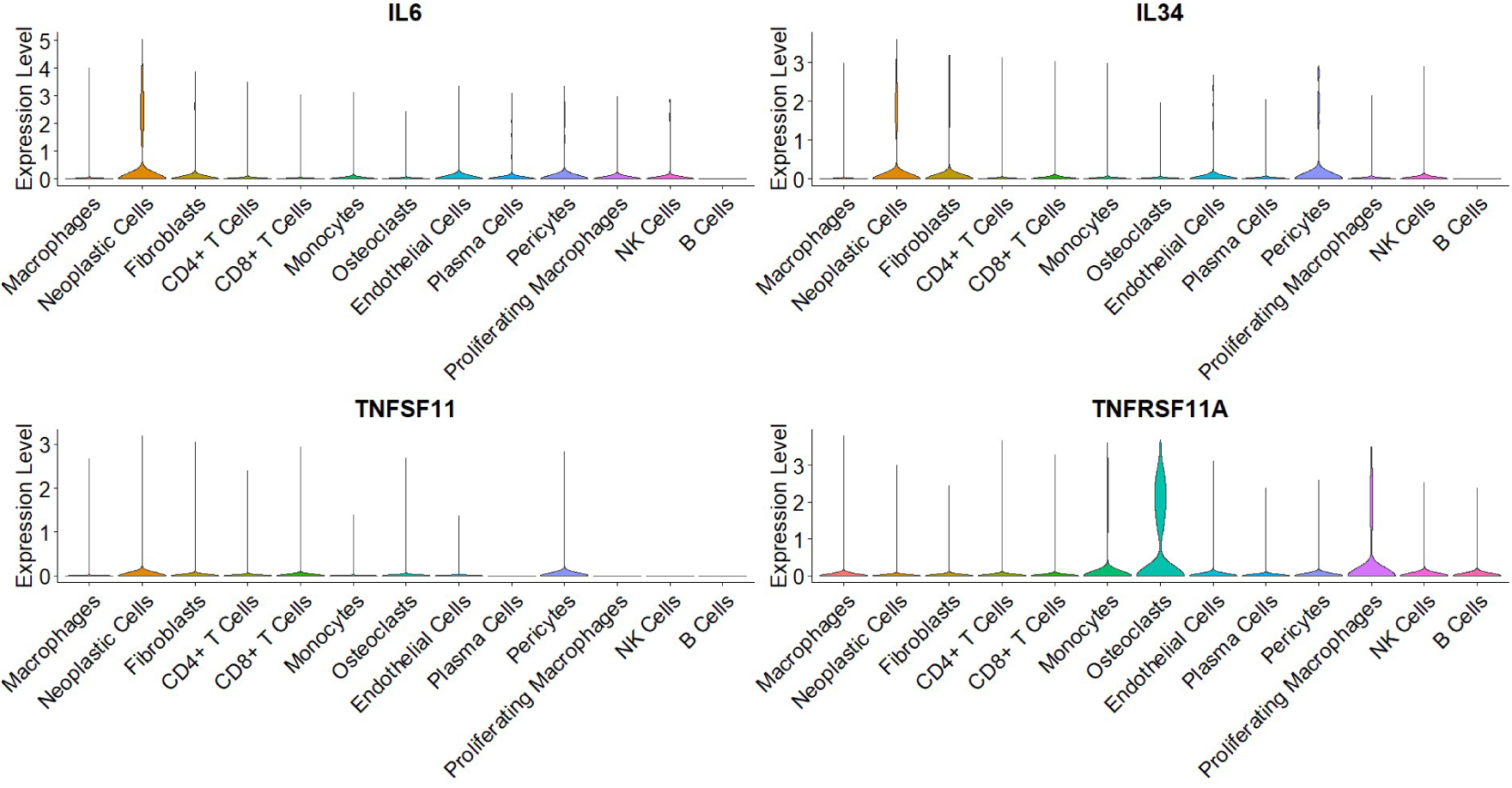
Expression of *IL6* and *IL34* in neoplastic cells, possibly binding CSF1R. While there is *TNFRSF11A* (RANK) expression in osteoclasts, there is no *TNFSF11* (RANKL) expression in any KPGCT cells.

**Supplemental Figure 10.**
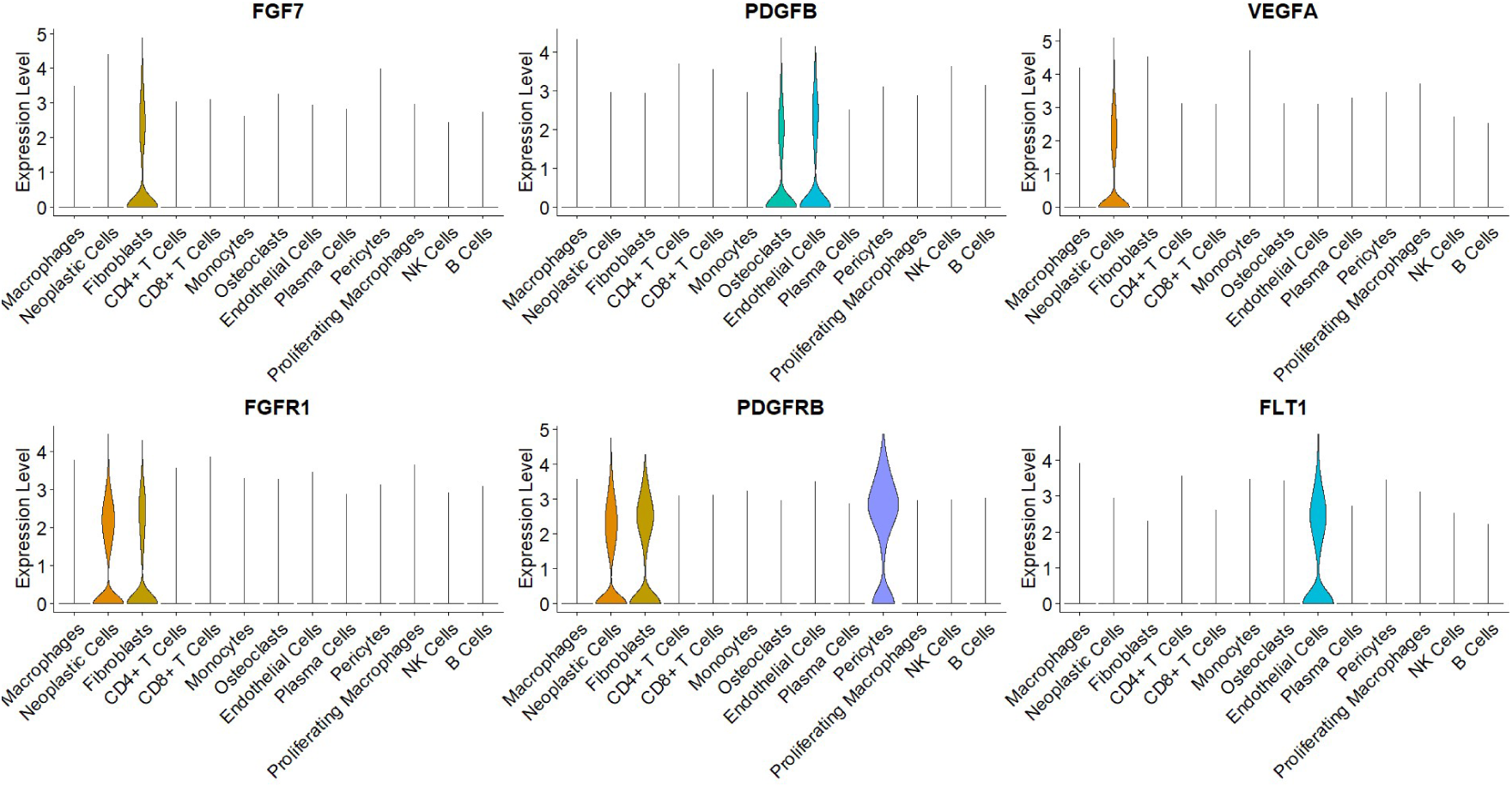
Expression of key signaling genes in the KPGCT microenvironment

**Supplemental Figure 11.**
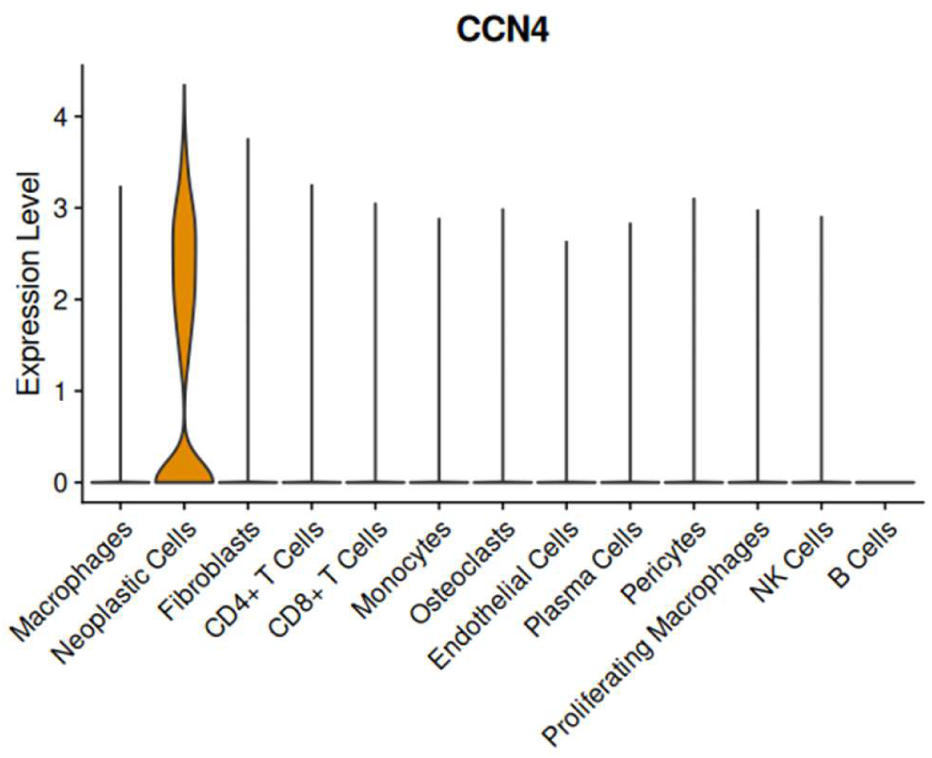
C*C*N4, a known activator of the hippo pathway, is highly expressed in neoplastic cells

**Supplemental Figure 12.**
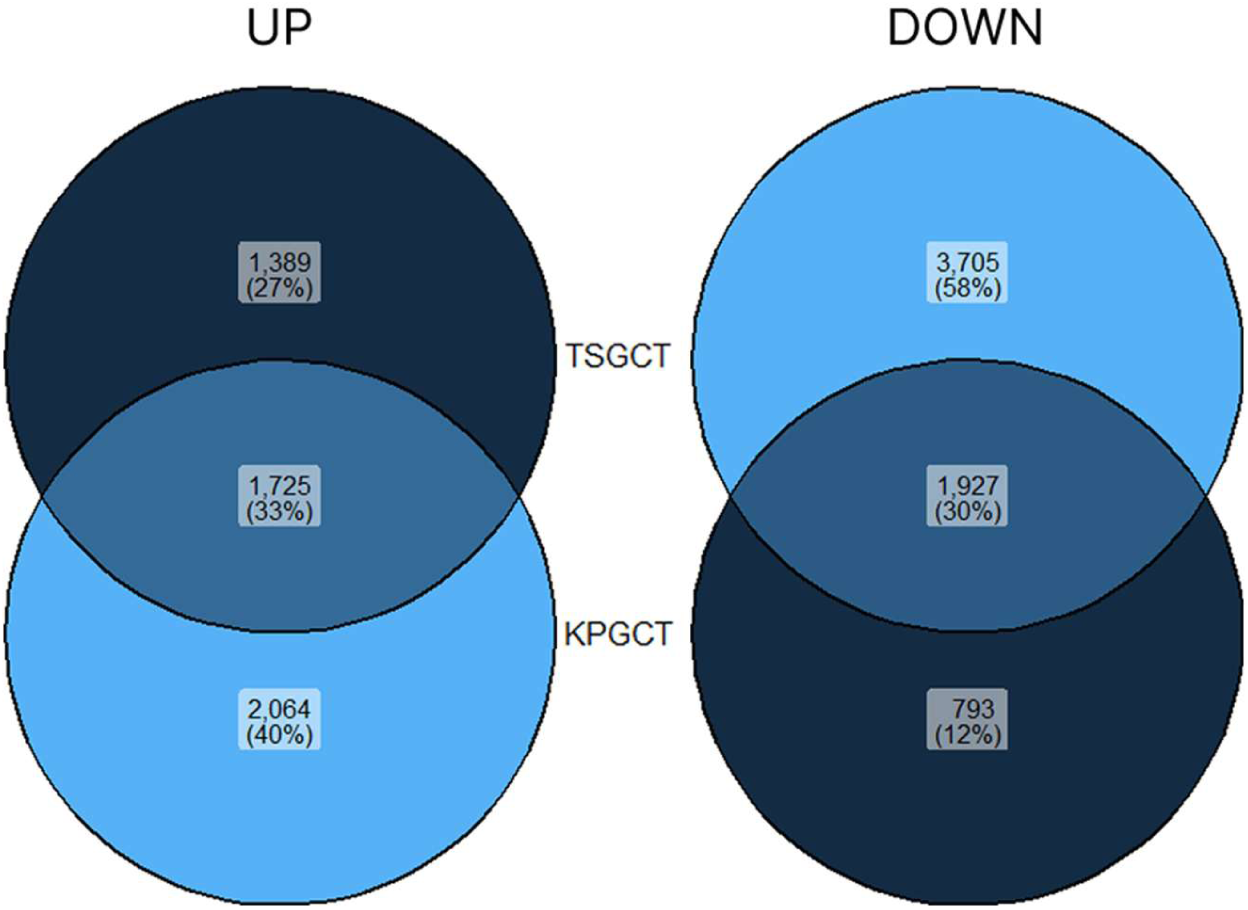
Overlap of differentially expression genes in TGCT and KPGCT

**Supplemental Figure 13.**
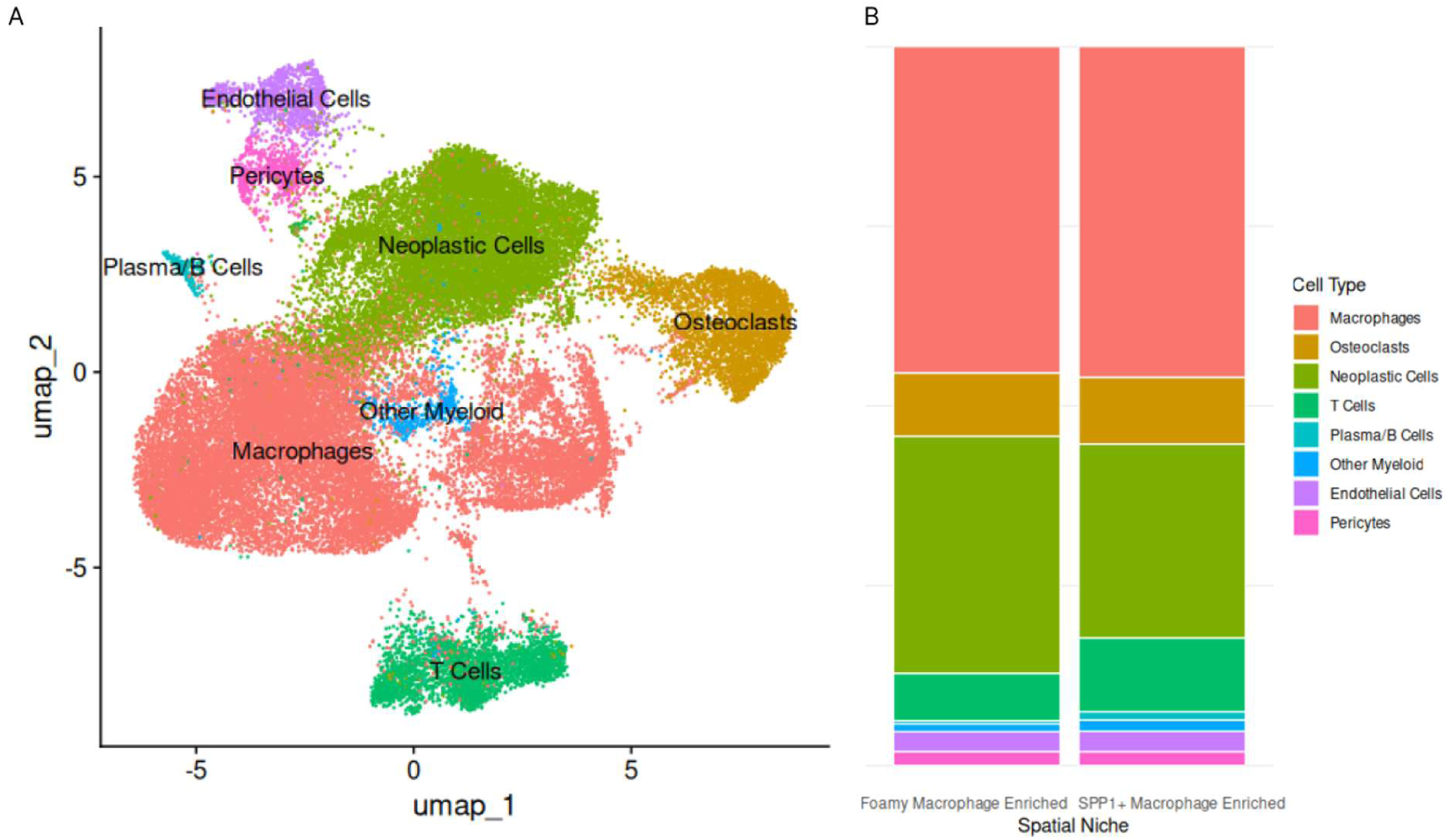
A) UMAP plot of cells sequenced through Xenium. B) Distribution of cells in different niches.

**Supplemental Figure 14.**
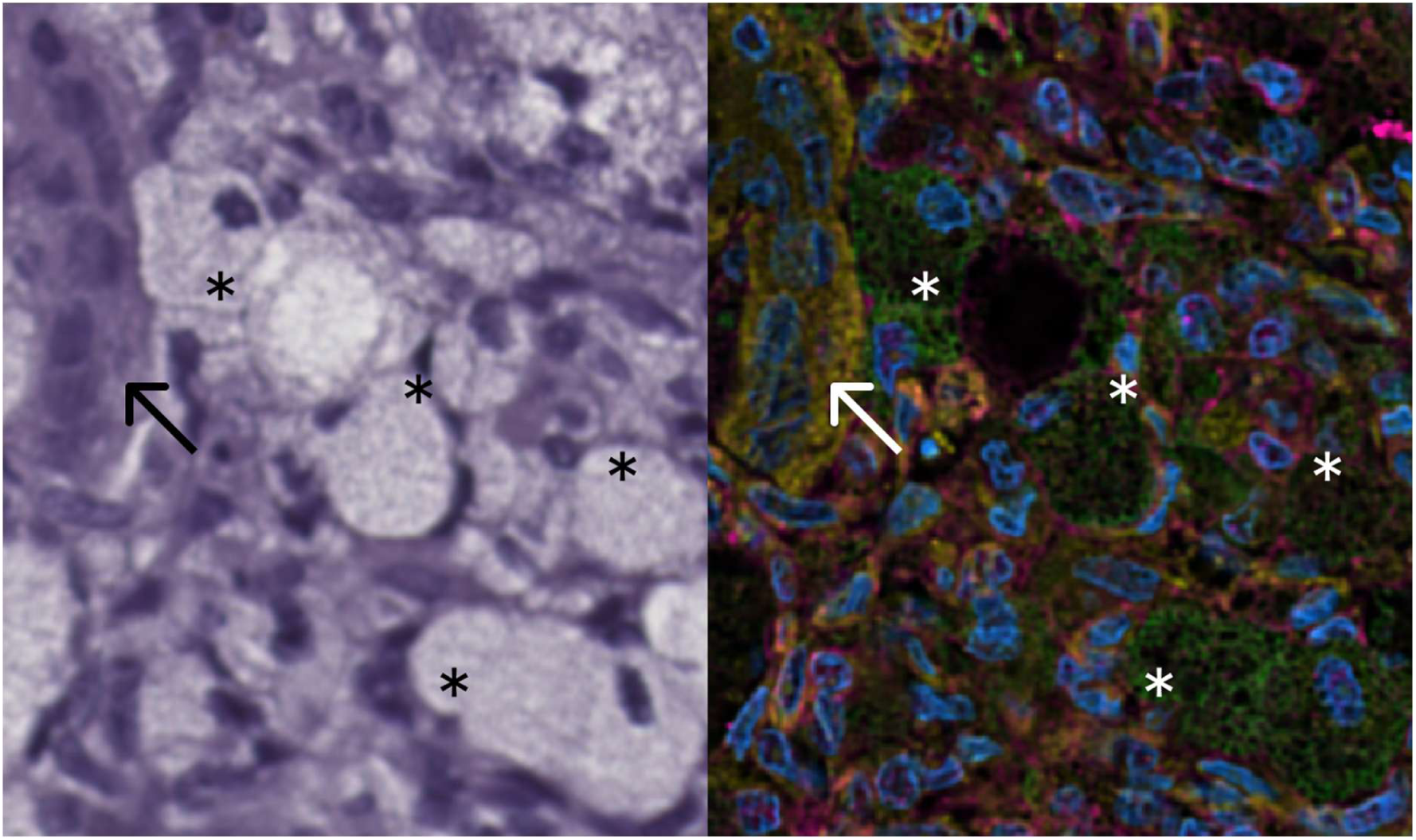
Foamy macrophages (asterisks) and osteoclast-like giant cells (arrows) are easily identified on both histology (left) and Xenium segmentation stains (right).

**Supplemental Figure 15.**
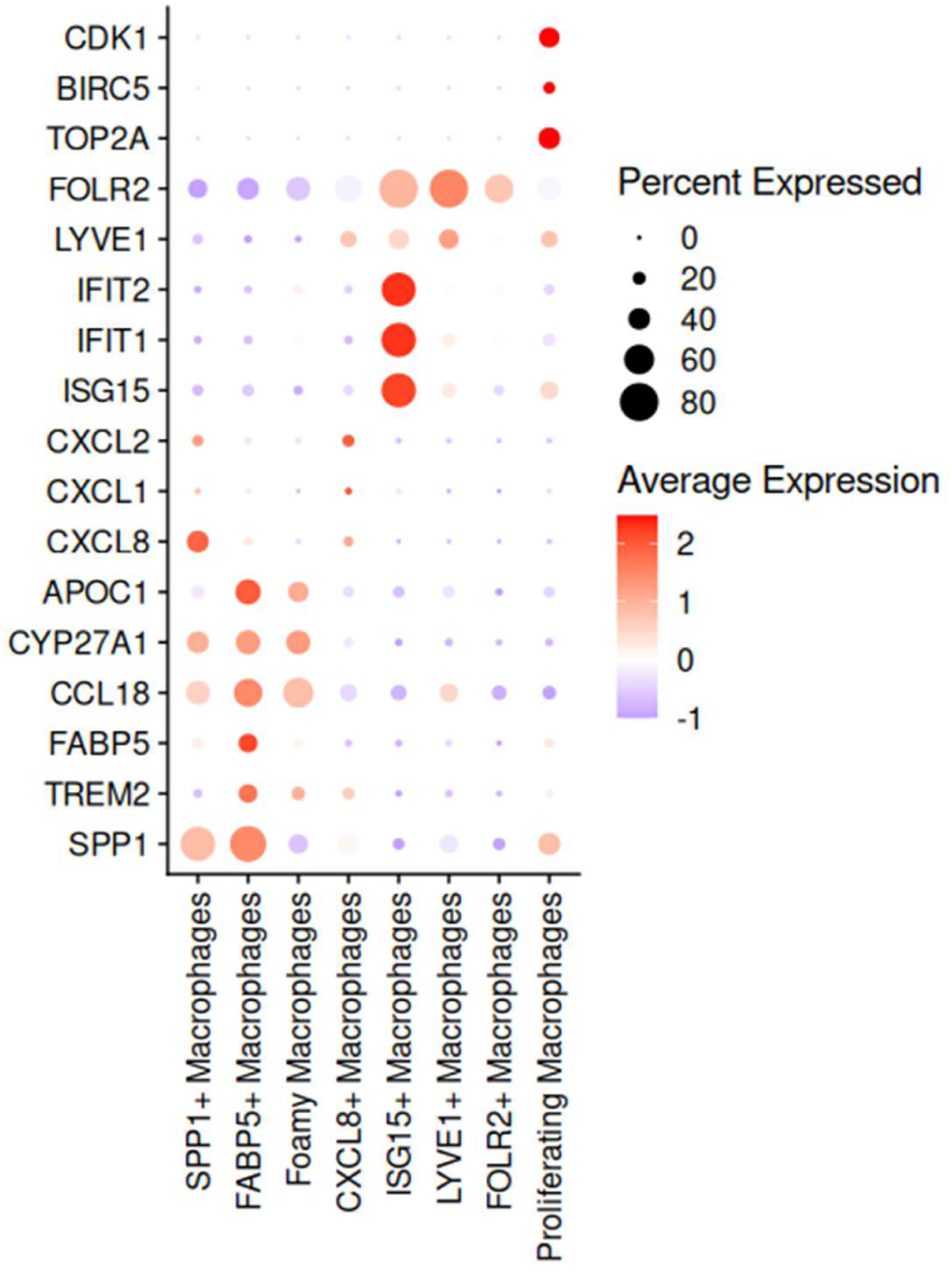
KPGCT macrophage subsets found in Xenium spatial transcriptomics are also found in snRNA-seq

**Supplemental Figure 16.**
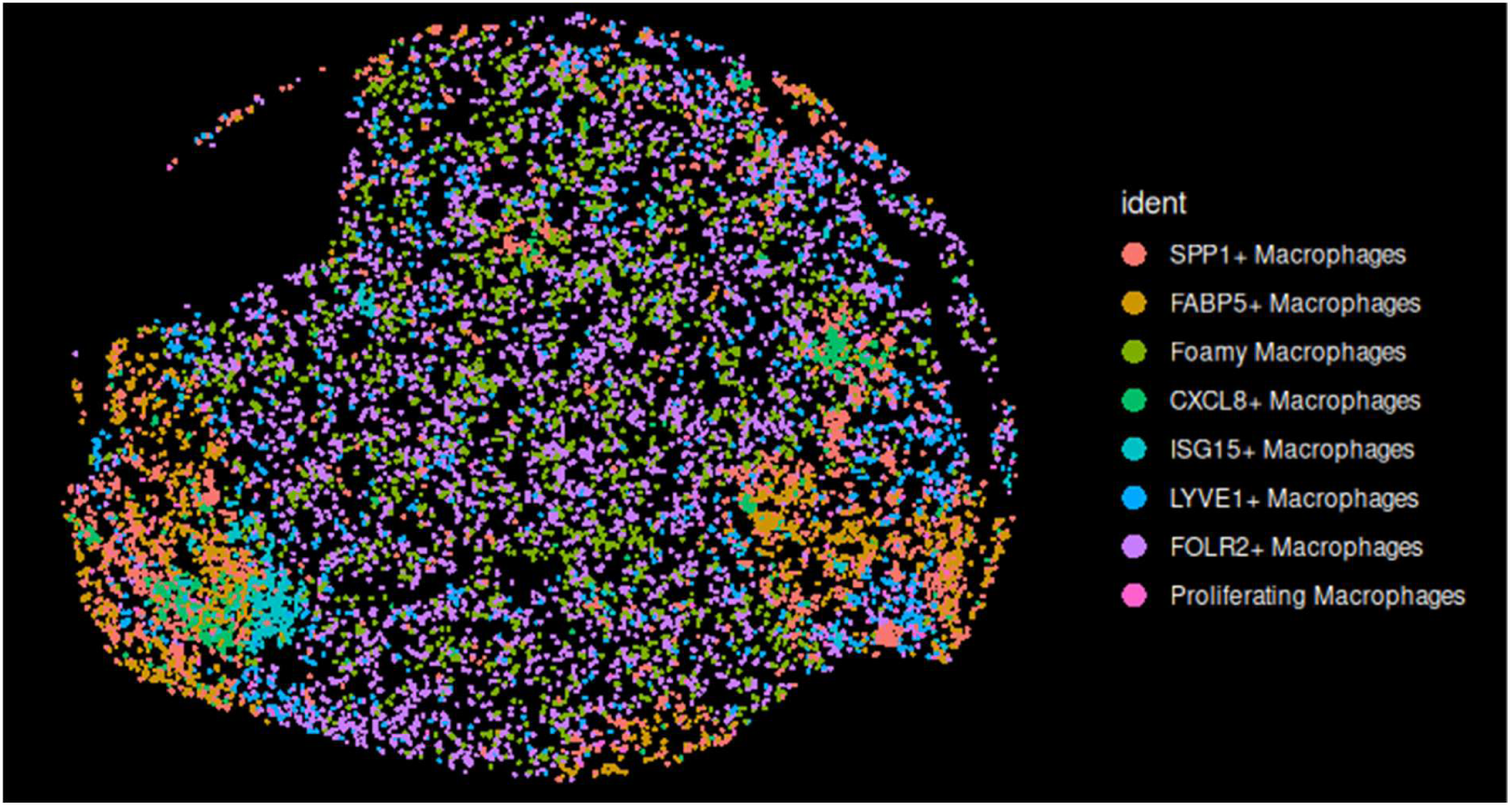
SPP1+, FABP5+, CXCL8+ and ISG15+ macrophages form in small areas, while foamy and FOLR2+ macrophages are more interspersed.

## References

1. K. J. Fritchie, et al., Xanthogranulomatous epithelial tumor: report of 6 cases of a novel, potentially deceptive lesion with a predilection for young women. Mod. Pathol. 33, 1889–1895 (2020).

2. C. A. Dehner, et al., Xanthogranulomatous epithelial tumors and keratin-positive giant cell-rich soft tissue tumors: two aspects of a single entity with frequent *HMGA2-NCOR2* fusions. Mod. Pathol. 35, 1656–1666 (2022).

3. P. Rungsiprakarn, et al., Keratin-Positive Giant Cell Tumor of Bone and Soft Tissue With HMGA2::NCOR2 Fusion in Children Under 10 With Response to Imatinib Therapy: A Case Series. *JCO Precis*. Oncol. 8, e2300659 (2024).

4. R. D. Whaley, et al., Xanthogranulomatous epithelial tumors/keratin-positive giant cell–rich tumors involving the head and neck: report of seven cases and review of the literature. Virchows Arch. 485, 605–613 (2024).

5. A. Agaimy, et al., Recurrent novel *HMGA2-NCOR2* fusions characterize a subset of keratin-positive giant cell-rich soft tissue tumors. Mod. Pathol. 34, 1507–1520 (2021).

6. A. R. Shih, et al., Clinicopathologic characteristics of poorly differentiated chordoma. Mod. Pathol. 31, 1237–1245 (2018).

7. S. D. Humble, V. G. Prieto, M. G. Horenstein, Cytokeratin 7 and 20 expression in epithelioid sarcoma. J. Cutan. Pathol. 30, 242–246 (2003).

8. J. Iwata, C. D. Fletcher, Immunohistochemical detection of cytokeratin and epithelial membrane antigen in leiomyosarcoma: a systematic study of 100 cases. Pathol. Int. 50, 7–14 (2000).

9. M. Miettinen, J. Limon, A. Niezabitowski, J. Lasota, Patterns of keratin polypeptides in 110 biphasic, monophasic, and poorly differentiated synovial sarcomas. Virchows Arch. Int. J. Pathol. 437, 275–283 (2000).

10. C. A. Dehner, D. Buehler, C. Hofich, K. C. Halling, A. L. Folpe, Novel HMGA2::COL14A1 Fusion Identified in Xanthogranulomatous Epithelial Tumor/Keratin-Positive Giant Cell Tumor. Genes. Chromosomes Cancer 63, e70010 (2024).

11. C. A. Dehner, et al., CSF1 expression in xanthogranulomatous epithelial tumor/keratin-positive giant cell-rich tumor. Hum. Pathol. 143, 1–4 (2024).

12. R. Perret, et al., Giant Cell Tumors With HMGA2::NCOR2 Fusion: Clinicopathologic, Molecular, and Epigenetic Study of a Distinct Entity. Am. J. Surg. Pathol. 47, 801 (2023).

13. M. Brahmi, et al., Complete response to CSF1R inhibitor in a translocation variant of teno-synovial giant cell tumor without genomic alteration of the CSF1 gene. Ann. Oncol. Off. J. Eur. Soc. Med. Oncol. 29, 1488–1489 (2018).

14. Y. Hao, et al., Dictionary learning for integrative, multimodal and scalable single-cell analysis. Nat. Biotechnol. 42, 293–304 (2024).

15. O. Franzén, L.-M. Gan, J. L. M. Björkegren, PanglaoDB: a web server for exploration of mouse and human single-cell RNA sequencing data. Database 2019, baz046 (2019).

16. M. I. Love, W. Huber, S. Anders, Moderated estimation of fold change and dispersion for RNA-seq data with DESeq2. Genome Biol. 15, 550 (2014).

17. V. Singhal, et al., BANKSY unifies cell typing and tissue domain segmentation for scalable spatial omics data analysis. Nat. Genet. 56, 431–441 (2024).

18. S. Jin, M. V. Plikus, Q. Nie, CellChat for systematic analysis of cell–cell communication from single-cell transcriptomics. Nat. Protoc. 20, 180–219 (2025).

19. G. Korotkevich, et al., Fast gene set enrichment analysis. [Preprint] (2021). Available at: https://www.biorxiv.org/content/10.1101/060012v3 [Accessed 1 August 2025].

20. G. Yu, C.-H. Gao, enrichplot: Visualization of Functional Enrichment Result. 10.18129/B9.bioc.enrichplot.

21. W. Luo, C. Brouwer, Pathview: an R/Bioconductor package for pathway-based data integration and visualization. Bioinforma. Oxf. Engl. 29, 1830–1831 (2013).

22. J. Cao, et al., A human cell atlas of fetal gene expression. Science 370, eaba7721 (2020).

23. D. G. P. van IJzendoorn, et al., Interactions in CSF1-Driven Tenosynovial Giant Cell Tumors. Clin. Cancer Res. 28, 4934–4946 (2022).

24. T. Yamashita, et al., NF-kappaB p50 and p52 regulate receptor activator of NF-kappaB ligand (RANKL) and tumor necrosis factor-induced osteoclast precursor differentiation by activating c-Fos and NFATc1. J. Biol. Chem. 282, 18245–18253 (2007).

25. Y.-N. Wang, W.-C. Chang, Induction of disease-associated keratin 16 gene expression by epidermal growth factor is regulated through cooperation of transcription factors Sp1 and c-Jun. J. Biol. Chem. 278, 45848–45857 (2003).

26. S. Ma, L. Rao, I. M. Freedberg, M. Blumenberg, Transcriptional control of K5, K6, K14, and K17 keratin genes by AP-1 and NF-kappaB family members. Gene Expr. 6, 361–370 (1997).

27. J. M. Boland, A. L. Folpe, J. L. Hornick, K. L. Grogg, Clusterin is Expressed in Normal Synoviocytes and in Tenosynovial Giant Cell Tumors of Localized and Diffuse Types: Diagnostic and Histogenetic Implications. Am. J. Surg. Pathol. 33, 1225 (2009).

28. K. Maly, et al., The Expression of Thrombospondin-4 Correlates with Disease Severity in Osteoarthritic Knee Cartilage. Int. J. Mol. Sci. 20, 447 (2019).

29. G. D. Jay, D. E. Britt, C. J. Cha, Lubricin is a product of megakaryocyte stimulating factor gene expression by human synovial fibroblasts. J. Rheumatol. 27, 594–600 (2000).

30. M. Matusiak, et al., Spatially Segregated Macrophage Populations Predict Distinct Outcomes in Colon Cancer. Cancer Discov. 14, 1418–1439 (2024).

31. P. Rogalla, et al., HMGI-C expression patterns in human tissues. Implications for the genesis of frequent mesenchymal tumors. Am. J. Pathol. 149, 775–779 (1996).

32. N. Dreux, et al., Value and limitation of immunohistochemical expression of HMGA2 in mesenchymal tumors: about a series of 1052 cases. Mod. Pathol. Off. J. U. S. Can. Acad. Pathol. Inc 23, 1657–1666 (2010).

33. J. S. A. Chrisinger, C. A. Dehner, Keratin-positive giant cell-rich tumor review. Semin. Diagn. Pathol. 42, 150939 (2025).

34. S. Sugita, et al., Diagnostic utility of CSF1 immunohistochemistry in tenosynovial giant cell tumor for differentiating from giant cell-rich tumors and tumor-like lesions of bone and soft tissue. Diagn. Pathol. 17, 88 (2022).

35. J. J. Thangaiah, J. W. Koepplin, A. L. Folpe, RNAscope CSF1 chromogenic in situ hybridization: a potentially useful tool in the differential diagnosis of tenosynovial giant cell tumors. Hum. Pathol. 115, 1–9 (2021).

36. T. Gauduchon, et al., Expanding the molecular spectrum of tenosynovial giant cell tumors. Front. Oncol. 12 (2022).

37. R. Felix, M. G. Cecchini, H. Fleisch, Macrophage Colony Stimulating Factor Restores in vivo Bone Resorption in the op/op Osteopetrotic Mouse. Endocrinology 127, 2592–2594 (1990).

38. S. Tanaka, et al., Macrophage colony-stimulating factor is indispensable for both proliferation and differentiation of osteoclast progenitors. J. Clin. Invest. 91, 257–263 (1993).

39. H. Yoshida, et al., The murine mutation osteopetrosis is in the coding region of the macrophage colony stimulating factor gene. Nature 345, 442–444 (1990).

40. B. R. MacDonald, et al., Effects of human recombinant CSF-GM and highly purified CSF-1 on the formation of multinucleated cells with osteoclast characteristics in long-term bone marrow cultures. J. Bone Miner. Res. 1, 227–233 (1986).

41. J.-Y. Blay, H. El Sayadi, P. Thiesse, J. Garret, I. Ray-Coquard, Complete response to imatinib in relapsing pigmented villonodular synovitis/tenosynovial giant cell tumor (PVNS/TGCT). Ann. Oncol. Off. J. Eur. Soc. Med. Oncol. 19, 821–822 (2008).

42. H. Gelderblom, et al., Vimseltinib versus placebo for tenosynovial giant cell tumour (MOTION): a multicentre, randomised, double-blind, placebo-controlled, phase 3 trial. The Lancet 403, 2709–2719 (2024).

43. Pimicotinib Significantly Improved Outcomes for Patients with TGCT in Phase III Trial. Available at: https://www.emdgroup.com/en/news/pimicotinib-topline-results-12-11-2024.html?global_redirect=1 [Accessed 28 May 2025].

44. W. D. Tap, et al., Pexidartinib versus placebo for advanced tenosynovial giant cell tumour (ENLIVEN): a randomised phase 3 trial. The Lancet 394, 478–487 (2019).

45. K. K. Sankhala, et al., A phase I/II dose escalation and expansion study of cabiralizumab (cabira; FPA-008), an anti-CSF1R antibody, in tenosynovial giant cell tumor (TGCT, diffuse pigmented villonodular synovitis D-PVNS). J. Clin. Oncol. 35, 11078–11078 (2017).

46. S. Roux, et al., RANK (Receptor Activator of Nuclear Factor kappa B) and RANK Ligand Are Expressed in Giant Cell Tumors of Bone. Am. J. Clin. Pathol. 117, 210–216 (2002).

47. T. Dong, et al., WISP1 inhibition of YAP phosphorylation drives breast cancer growth and chemoresistance via TEAD4 activation. Anticancer. Drugs 36, 157–176 (2025).

48. A. Vassilev, K. J. Kaneko, H. Shu, Y. Zhao, M. L. DePamphilis, TEAD/TEF transcription factors utilize the activation domain of YAP65, a Src/Yes-associated protein localized in the cytoplasm. Genes Dev. 15, 1229–1241 (2001).

49. T. A. Yap, et al., YAP/TEAD inhibitor VT3989 in solid tumors: a phase 1/2 trial. Nat. Med. 1–10 (2025). 10.1038/s41591-025-04029-3.

50. F. Medeiros, et al., Frequency and characterization of HMGA2 and HMGA1 rearrangements in mesenchymal tumors of the lower genital tract. Genes. Chromosomes Cancer 46, 981–990 (2007).

51. E. F. P. M. Schoenmakers, et al., Recurrent rearrangements in the high mobility group protein gene, HMGI-C, in benign mesenchymal tumours. Nat. Genet. 10, 436–444 (1995).

52. U. Unachukwu, K. Chada, J. D’Armiento, High Mobility Group AT-Hook 2 (HMGA2) Oncogenicity in Mesenchymal and Epithelial Neoplasia. Int. J. Mol. Sci. 21, 3151 (2020).

53. A. Dahlén, et al., Fusion, Disruption, and Expression of HMGA2 in Bone and Soft Tissue Chondromas. Mod. Pathol. 16, 1132–1140 (2003).

54. H. Bartuma, et al., Assessment of the clinical and molecular impact of different cytogenetic subgroups in a series of 272 lipomas with abnormal karyotype. Genes. Chromosomes Cancer 46, 594–606 (2007).

55. A. Agaimy, et al., HMGA2-WIF1 Rearrangements Characterize a Distinctive Subset of Salivary Pleomorphic Adenomas With Prominent Trabecular (Canalicular Adenoma-like) Morphology. Am. J. Surg. Pathol. 46, 190–199 (2022).

